# Widespread roles of *Trypanosoma brucei* ATR in nuclear genome function and transmission are linked to R-loops

**DOI:** 10.1101/2021.09.09.459654

**Authors:** J.A. Black, K. Crouch, E. Briggs, L. Lemgruber, C. Lapsely, L. R. O. Tosi, J. C. Mottram, R. McCulloch

## Abstract

Inheritance of aberrant chromosomes can compromise genome integrity and affect cellular fitness. In eukaryotes, surveillance pathways and cell cycle checkpoints monitor for aberrant DNA transmission and the ATR kinase, a regulator of the DNA damage response, plays a pivotal role. Prior work revealed that ATR acts during antigenic variation in *Trypanosoma brucei* mammal-infective life cycle forms and that its loss is lethal, but how widely ATR operates in genome maintenance is largely unknown. Here, we show that after prolonged ATR depletion by RNAi *T. brucei* continues to synthesise DNA and enters new rounds of cell division, despite increased genome damage. Furthermore, we detect defective chromosome segregation, ‘micronuclei’ formation and disruption of the nuclear architecture. RNA-seq revealed that loss of ATR affects the expression of nearly half the genes in the genome, including both RNA Polymerase I and II transcription. Using ChIP-seq of yH2A and DRIP-seq, we reveal overlapping signals for genome damage and R-loops after ATR depletion in all intergenic regions. In addition, we report reduced R-loop levels and accumulation of yH2A signal within centromeres. Together, our data indicates widespread roles of ATR in *T. brucei*, including differing roles in R-loop homeostasis during multigene transcription and in chromosome segregation.

## Introduction

Once per cell cycle, the nuclear genome is duplicated and physically segregated, with each daughter cell normally inheriting a faithful copy^1^. However, during cell growth and genome replication the DNA template can become damaged, aberrant DNA structures may form or chromosomes may fail to segregate correctly^2,3^, leading to imbalances in gene expression and genome instability. To ensure high fidelity genome inheritance, eukaryotes use a range of surveillance pathways and cell cycle checkpoints as controls to limit cell cycle progression in the presence of aberrant DNA content, safeguarding the genome. Cell cycle checkpoints^2,4^ act at boundaries between cell cycle stages as well as within stages, such as the intra-S phase checkpoint, which monitors DNA replication (reviewed by^5^), and the spindle assembly checkpoint (SAC), which monitors chromosome segregation (reviewed by^6^). Protein kinases dictate the activity of such cell cycle checkpoints.

Genome surveillance involves the DNA Damage Response (DDR), an interconnected network of repair pathways that intersect with cell cycle checkpoints (reviewed by^2,7^). At the apex of the eukaryotic DDR sit members of an atypical Phosphatidylinositol 3-kinase (PI3K)-like protein kinase family^8^, acting to recognise and signal the presence of genome damage, thereby enacting and orchestrating the appropriate repair pathway. One such PI3K-like kinase is called ataxia-telangiectasia and Rad3 related, or ATR (reviewed by^8^). The function of ATR during S-phase of the cell cycle has been extensively characterised. ATR, when complexed with an obligatory binding partner (ATRIP^9^, and ETAA1^10^ in vertebrates), is activated by the accumulation of single-stranded DNA (ssDNA), a hallmark of DNA replication stress during S phase. The ssDNA is coated and stabilised by the Replication Protein A (RPA) complex and the RPA-ssDNA structure can partially activate ATR^11^. Independent loading of the heterotrimeric 9-1-1 complex and TopBP1^12–15^ onto DNA promotes full activation of ATR, leading to phosphorylation of the Checkpoint 1 (CHK1) kinase (reviewed by^16^). CHK1 activation prevents cell cycle progression prior to mitosis, ultimately through inhibition of CDC25A^17^. One aspect of ATR’s replication-associated functions is to tackle RNA-DNA hybrids (R-loops) that form during clashes with transcription^18–20^. In addition to S-phase functions, ATR acts during S/G_2_^4^ and G_2_/M cell cycle transitions^21,22^, and has been implicated in recognising stalled RNA Polymerase (Pol) II^23^, where it acts through a phosphatase complex that has been shown to act in transcription termination in *Trypanosoma brucei*^24,25^. More recently, ATR was shown to ensure faithful chromosome segregation during mitosis^26^. Here, ATR activation by RPA accumulation at centromeres occurs due to the formation of RNA Pol II-derived R-loops. Once activated, ATR directs chromosome segregation by initiating an aurora kinase B (AUKB)-dependent pathway. This function appears to be distinct from more canonical ATR repair activities^26^. Finally, ATR has been described as having widespread cellular roles in protecting the integrity of the nuclear envelope, nucleolus and centrosome (reviewed in ^27^ and^28–31^).

*T. brucei* is a protozoan parasite and one aetiological agent of debilitating and often fatal trypanosomiases diseases that afflict humans and animals worldwide^32^. A *T. brucei* homologue of ATR (TbATR) has been identified^33^ and its depletion through RNAi in mammal-infective (bloodstream form, BSF) cells shown to lead rapidly to changes in the control of RNA Pol I-transcribed Variant Surface Glycoprotein (VSG) genes that are expressed monoallelically and undergo switching in expression to mediate host immune evasion^34^. TbATR depletion in BSF cells is lethal and associated with cell cycle progression defects^35,36^ and accumulation of phosphorylated histone H2A (yH2A), indicating genome damage^34^. These data indicate wider, essential roles for ATR in African trypanosomes whose basis have not been dissected. Surprisingly, in tsetse-infective (procyclic form, PCF) *T. brucei* cells RNAi depletion or chemical inhibition of ATR appears not to affect growth, though cell cycle progression and DNA double-strand break repair are impaired^37^. Although many pathways dictating genome repair and cell cycle progression are conserved in *T. brucei*, these parasites lack identifiable homologs of ATR pathway members, including CHK1 and Cdc25^38^ and in the related kinetoplastid parasite *Leishmania*, divergent 9-1-1 complex functions have been reported^39,40^, suggesting the kinetoplastid ATR pathway may operate distinctly from that of a ‘model’ eukaryote. Thus, what roles TbATR may serve in *T. brucei*, outwith immune evasion, are largely unknown, and how TbATR contributes to cell growth and survival is unclear. To answer these questions, we tested for roles of TbATR in monitoring cell cycle progression, maintaining genome stability, directing genome transmission, and protecting nuclear structure in BSF *T. brucei* parasites. We show that prolonged loss of TbATR is associated with chromosome segregation defects and genome-wide damage accumulation. We reveal, by RNA-seq analysis, significant alteration of transcription across the genome and demonstrate that this perturbation is linked to the distribution of R-loops, including during RNA Pol I and Pol II transcription, and at centromeres. Our work indicates TbATR monitors R-loops genome-wide, including during chromosome segregation, and thereby reveals functions for ATR during transcription and mitosis in the African trypanosome.

## Materials and Methods

Information regarding reagents is listed in Table S2.

### Oligonucleotide Sequences

All oligonucleotide sequences are shown in Table S2. All were designed using CLC Genomics Workbench 7 (Qiagen) and synthesized by Eurofins Genomics (http://www.eurofins.com). All sequence information used was taken from TriTrypDB (http://tritrypdb.org/tritrypdb) and analysed *in silico* using BLAST (NCBI)^41^ using default parameters.

### Plasmid Design and Cloning

To endogenously tag TbATR at the N-terminus, the vector pEnT6B (modified from^42^) was used. The construct (termed ^12myc^TbATR) was transformed into WT427 cells lacking one allele of the TbATR kinase which was replaced with the neomycin (NEO) resistance gene to confer resistance to G418 (Neomycin). One endogenous allele was replaced using a modified pmtl23 vector as described^43^.

### Polymerase Chain Reaction

Genomic DNA for PCR analysis was extracted using the Qiagen Blood and Tissue Extraction Kit^®^ (Qiagen) as per the manufacturer’s instructions. PCR confirmation of endogenously tagged ATR cell lines was performed using NEB Taq DNA polymerase (NEB) as per the manufacturer’s instructions. For construction of plasmids, Phusion^®^ High-Fidelity DNA Polymerase (NEB) was used as per manufacturer’s instruction.

### Trypanosome Culture

All cell lines used in this study are derived from *Trypanosoma brucei brucei* Lister 427^44^ cells. TbATR RNAi cell lines are as described ^34,35,45^ and 2T1 cells^46^ were used as control cells and maintained as described. BSF RNAi cell lines were cultured in HMI-9 medium ^47^ supplemented with 10% (v/v) low-tetracycline foetal calf serum (FCS; Gibco) at 37 °C in 5% C0_2_ and maintained in 5 µg.mL^-1^ Hygromycin (InvivoGen) and 5 µg.mL^-1^ Phleomycin (InvivoGen).

Endogenously tagged cell lines were maintained in HMI-9 supplemented with 20 % (v/v) FCS at 37 °C in 5% C0_2_ and maintained in 10 µg.mL^-1^ Blasticidin (InvivoGen). Parasite density was monitored using a haemocytometer.

### Parasite Growth and Cell Cycle Analysis

For growth analysis, cells were set at 1 x10^4^ cells/mL in HMI-9 containing the appropriate selective antibiotics. Cells were grown in the presence or absence of methyl methanesulfonate (MMS; Sigma) at a concentration of 0.0003% for 72hrs. Population density was assessed every 24hrs. MMS was stocks were diluted in HMI-9.

### Immunoblotting Analysis

Immunoblotting to detect myc-tagged ATR was performed as follows. Approximately 2.5 x10^6^ cells were harvested by centrifugation (1620 x *g* for 10 mins), the pellet washed in 1x PBS then re-suspended in 1x protein loading buffer (4x NuPAGE^®^ LDS sample buffer [Invitrogen], 1x PBS and 25 uL ß-mercaptoethanol, 2x Roche cOmplete Mini protease inhibitor cocktail tablets) and denatured immediately at 100 °C for 10 mins. Whole cell lysates were separated by SDS-PAGE on 3-8% Tris acetate NuPAGE^®^ Novex^®^ pre-cast gels and run as per manufacturer’s instructions. For blotting, proteins were transferred onto a PVDF membrane (Amersham Bio) at 400 mA for 1 hr followed by 90 mA overnight at 4 °C. Transfer buffer used: 25 mM Tris pH 8.3, 192 mM Glycine, 20 % [v/v] methanol. The membrane was washed in 1x PBST (1x PBS, 0.01 % Tween-20 [Sigma]) for 10 mins and incubated in blocking solution (1x PBST, 5 % non-fat Milk powder [Marvel]) for 1 hr. After, the membrane was washed for 10 mins in 1x PBST then incubated with the primary antibody (mouse anti-myc clone 4A6 [Milipore]; 1:7000) dilute in blocking solution. Mouse anti-EF1 alpha (Millipore; 1:20 000) was used as a loading control. After, the membrane was washed 2x for 10 mins each in 1x PBST then incubated in blocking solution containing goat anti-mouse HRP conjugate (Thermo; 1: 3000). The membrane was subsequently washed for 3x in 1x PBST then incubated for 5 mins in the dark with ECL Primer Western Blotting Detection Reagent (Amersham) and exposed onto ECL Hyperfilm (Amersham).

### Analysis of Replicating DNA by EdU Incorporation and yH2A Detection

Cells were set at a concentration of 1 x10^5^ cells/mL 24 hrs prior to EdU addition (5-ethynl-2’-deoxyuridine; Life Technology, Thermo Scientific) in media free from thymidine prepared using Iscove’s Modified Dulbecco’s Medium (IMDM; Gibco), 10 % [v/v] FBS (Gibco), HMI mix (0.05 mM bathocuproine disulfonic acid, 1 mM sodium pyruvate and 1.5 mM L-cysteine [Sigma]), 1 mM hypoxanthine (Sigma) and 0.00014 % 2-mercaptoethanol (Sigma). EdU labelling was performed exactly as described^48^. After 4 hrs, cells were harvested by centrifugation and fixed in 4% FA (Methanol free) onto glass slides (ThermoFisher). The wells were washed in 1x PBS and then treated with 100 mM Glycine for 20 mins. After, the cells were permeabilised in 0.2% Triton X-100 (Sigma) in 1x PBS for 10 mins then washed as before. The wells were subsequently blocked for 1 hr in blocking solution (25 µL of 1% BSA [Sigma], 0.2% Tween-20 in 1x PBS) then incubated with 25 µL of the Click-iT EdU detection mix (prepared as per manufacturer’s instructions using Alex Fluor 455^®^; Life Technologies, ThermoFisher) for 1 hr in the dark. The wells were then washed in 1x PBS three times. Immunofluorescence for yH2A was then performed. Wells were washed in 1x PBS then incubated with anti yH2A 1:1000 (In house; diluted in blocking solution) for 1 hr. After, wells were washed as before and incubated for 1 hr with the secondary antibody Alexa Fluor^®^ 488 anti-rabbit 1:1000 (ThermoFisher). Next, wells were washed then incubated for 5 mins in DAPI Fluoromount G (Southern Biotech) to stain nuclear and kinetoplastid DNA. Slides were stored at 4 °C until required.

### Transmission Electron Microscopy

For transmission electron microscopy, ∼ 5 x10^6^ cells were fixed in 2.5% glutaraldehyde and 4% paraformaldehyde (PFA) in 0.1 M sodium cacodylate buffer (pH 7.2). Samples were then post-fixed for 1h in 1% osmium tetroxide in 0.1 M sodium cacodylate buffer in the dark. Samples were washed several times with 0.1 M sodium cacodylate buffer then stained (*en bloc*) with 0.5% aqueous uranyl acetate, then dehydrated in acetone solutions (30, 50, 70, 90 and 100%). Samples were embedded in epoxy resin and sectioned (ultrathin sectioning, 60 nm thick) then visualised on a Tecnai T20 transmission electron microscope (ThermoFisher) operating at 120 kV.

### Telomere Fluorescence *In Situ* Hybridisation (Telo-FISH)

Approximately 5 x10^6^ cells were collected, washed by centrifugation (3000 rpm for 3 mins) then fixed in 4% FA (Methanol free) for 4 mins. The cells were left to settle on a Poly-L-Lysine (Sigma) treated slide for 20 mins then subsequently washed in 1x PBS. Cells were permeabilised in 0.1% Triton X-100 (Sigma) in 1x PBS for 5 mins then washed as before. The slides were dehydrated for 5 mins each in pre-chilled ethanol in ascending concentrations (70-90-100 %). After, the slides were left to dry during which the telomere-hybridisation probe was prepared. The Telomere PNA FISH Kit/FITC (DAKO) was used to probe for telomeres. The telomere probe was heated to 85 °C (water bath) in hybridisation solution (50% Formamid, 10% Dextran, 2x SSPE Buffer [1x SSPE: 0.18 M NaCl, 10 mM NaH_2_P0_4_, 1 mM EDTA, pH 7.7] buffer used at pH 7.9). Ten microliters of the probe were diluted in 60 µL hybridisation solution (per sample) and after heating, this solution was added to the slides, the slide sealed and further heated at 95 °C for 5 mins (water bath). The slide was then placed for 16 hrs in a 37 °C incubator. The slide was washed in 2x SSC (ThermoFisher)/50% Formamide for 30 mins at 37 °C then in 0.2x SSC for 60 mins at 50 °C and finally in 4x SSC for 10 mins at room temperature. The seal and supernatant were removed and the slides DAPI stained as above.

### Immunofluorescence Analysis

Immunofluorescence analysis to detect myc tagged ATR was performed exactly as described^34,45^. Anti-myc Alexa Fluor^®^ 488 conjugate (Millipore) at a concentration of 1:500 was used to detect myc tagged ATR and incubated at room temperature for 1 h in blocking buffer (1 % BSA [Sigma], 0.2 % Tween-20 in 1x PBS) in a wet chamber. For spindle staining, mouse anti-KMX-1 antiserum (1: 10; kind gift Dr Tansy Hammarton, University of Glasgow) was diluted in 1 % BSA only. Anti-mouse Alexa Fluor^®^ 594 (Thermo; 1:1000) was used to detect anti-KMX-1.

### Imaging and Image Processing

Images were captured on an Axioskop2 (Carl Zeiss, Germany) fluorescent microscope with a 63x lens and were acquired with ZEN Blue software (Carl Zeiss, Germany). For images captured on a DeltaVision Core microscope (AppliedPrecision, GE), a 1.40/100 x lens was used, and images were acquired using SoftWoRx suit 2.0 (AppliedPrecision, GE). Z-stacks were acquired and images deconvolved (conservative ratio; 1024×1024 resolution). ImageJ (Fiji; ^49^) was used to remove the background of images and for counting cells or quantifying fluorescence intensity. False colours were assigned to fluorescent channels. Brightness and contrast were set relative to the unstained controls. Signal was enhanced where appropriate to improve visualisation. Super-resolution structural illumination (3D-SIM) images were captured on an Elyra PS.1 Super Resolution microscope (Carl Zeiss, Germany) using ZEN Black Edition Imaging Software Tool suite (Carl Zeiss, Germany). A 1.4/63x lens was used to capture Z-stacks. 3D rendered images were produced using IMARIS software (v8.2; Oxford Instruments) from super-resolution images. Fluorescence intensity was quantified by measuring DAPI and myc signal within a region of interest (ROI) drawn around individual cell nuclei. The background was subtracted using a radius of 50 pixels. The mean intensity of the pixels was calculated in MS Excel (Microsoft^®^). Raw values are available in Table S1. Analysis was performed in ImageJ (Fiji)^50^.

### RNAseq Dataset

Raw sequence reads for RNAseq analysis were retrieved from the accession number provided here^34^ and analysed exactly as described^34^. TopGO was used to perform a GOterm analysis^51^.

### ChIPseq and DRIPseq Analysis

Normalised sequence reads for yH2A ChIPseq analysis were retrieved from the accession number provided^34^ and analysed as described. For DRIPseq, samples were prepared exactly as described^52^. Briefly, 2×10^8^ cells were harvested by centrifugation, fixed in 1 % formaldehyde (5 mins shaking; room temperature) then quenched in 125 mM Glycine. After, cells were re-suspended in Glycine Stop-Fix Solution (ChIP-IT Enzymatic Express Kit; Active Motif) and kept shaking for 5 mins at room temperature. Cells were lysed (as per manufacturer’s instructions) then chromatin extracted and digested as described using the Enzymatic Shearing Cocktail (37 °C) generating ∼ 200 bp sized fragments. R-Loops were immunoprecipitated using 4.5 ng of S9.6 antibody (Kerafast). RNase H1 treated samples were prepared as described^52^. Library preparation was conducted with a TruSeq ChIP Library Preparation Kit (Illumina) and 300 bp fragments (adaptors included) size selected using Agencourt AMPure XP beads (Beckman Coulter). Samples were subject to sequencing on an Illumina NextSeq 500 Platform. Sequencing was performed at Glasgow Polyomics. The resultant sequence reads were trimmed (using TrimGalore; default settings; Babraham Bioinformatics) and BowTie2^53^ used to align the sequences to the Lister 427 TbHGAPv10 genome^44^. Samples were filtered using SAMtools^54^ (MapQ value <1) and normalised to the respective input controls using DeepTools bamCompare (SES method – 50bp non-overlapping bins^55,56^) and the fold change expressed as a ratio of the reads. Sequence was examined in the Integrative Genome Browser (IGV^57^). Additional plots were generated in DeepTools (plotHeatmap; plotProfile) or in Gviz^58^. Sequence data was part analysed using the Galaxy Server (usegalaxy.org)^59^.

### Data Visualisation and Statistical Analysis

Data was visualised using Prism v9 (GraphPad) or in RStudio using ggplot2^60^ and Gviz^58^. ChIPseq and DRIPseq data were examined in IGV^57^. Sequence datasets were analyses as described in the appropriate methods section. All statistical analysis was in R or using Prism. Appropriate tests were selected and are as described in the corresponding figure legends or text. MS Excel (Microsoft^®^) was used to prepare data tables. Some sequence data analysis was performed using Galaxy server (usegalaxy.org^59^). All figures were assembled in and exported using BioRender.com.

### Data Availability

RNA-seq, yH2A ChIP-seq and DRIP-seq datasets have been deposited in the European Nucleotide Archive and can be accessed using the accession number: PRJEB23973^34^.

## Results

### ATR depletion in *T. brucei* compromises nuclear architecture

Previously, we^34^ and others^35^ documented that TbATR loss results in a transient G2/M cell cycle stall, manifesting as increased numbers of cells which have duplicated their kinetoplast but not nucleus. To ask whether loss of TbATR has wider nuclear effects, we first tested for compromised nuclear integrity in BSF cells by targeting TbATR using a tetracycline inducible RNAi system (as described in ^34,35,45^) and examining nuclear morphology by DAPI staining. Nuclei were scored in four categories: ‘no defect’ (where the nucleus looked comparable to a wild type (WT) cell nucleus); ‘fragmented’ (multiple, typically smaller nuclei were seen); ‘blebs’ (nuclear protrusions were seen, similar to effects observed after RNAi of nuclear laminar factors^61^); and ‘aberrant’ (nuclear morphology deviated from WT but no single, clear defect was obvious). We performed this analysis using two independently generated RNAi clones (CL1 and CL2), with cells collected after growth for 24, 36 and 48 hrs with and without induction of RNAi. Representative images of RNAi induced CL1 cells are shown in Fig.1A, and uninduced cells at 36 hrs are shown in Fig.S1A; quantification of the data is shown in Fig.1B. In uninduced cells, >95 % harboured nuclei with no observable defect at all time points. After RNAi induction the proportion of cells with abnormal nuclei increased from 24-48 hrs (∼5-40% in CL1; ∼20-60% in CL2). Most abnormal nuclei were classed as aberrant, possessing non-uniform deviations in size or shape. However, many examples were seen of RNAi induced cells that harboured nuclei with blebs, and a smaller but still clearly increased proportion of cells harboured fragmented nuclei, indicating a loss of nuclear integrity and/or uneven segregation of nuclear DNA content. A complication in scoring for fragmented nuclei is the potential to confuse DAPI staining of small nuclei with increased numbers of kinetoplasts. To begin to test this, and to further examine the underlying ultrastructure of the nucleus in TbATR depleted cells, we performed transmission electron microscopy 24 hrs (Fig.S1B) and 36 hrs (Fig.S1C) after RNAi. In keeping with the DAPI analysis, nuclear blebs and aberrant nuclear membranes were detected, in addition to cells with no clear nuclear defects, and cells with fragmented nuclei. Cells with multiple kinetoplasts were also detected, in keeping with replication of the kDNA being unperturbed after TbATR loss^34^. Together, these data indicate that prolonged depletion of TbATR compromises both nuclear ultrastructure and nuclear segregation in BSF cells.

**Figure 1.**
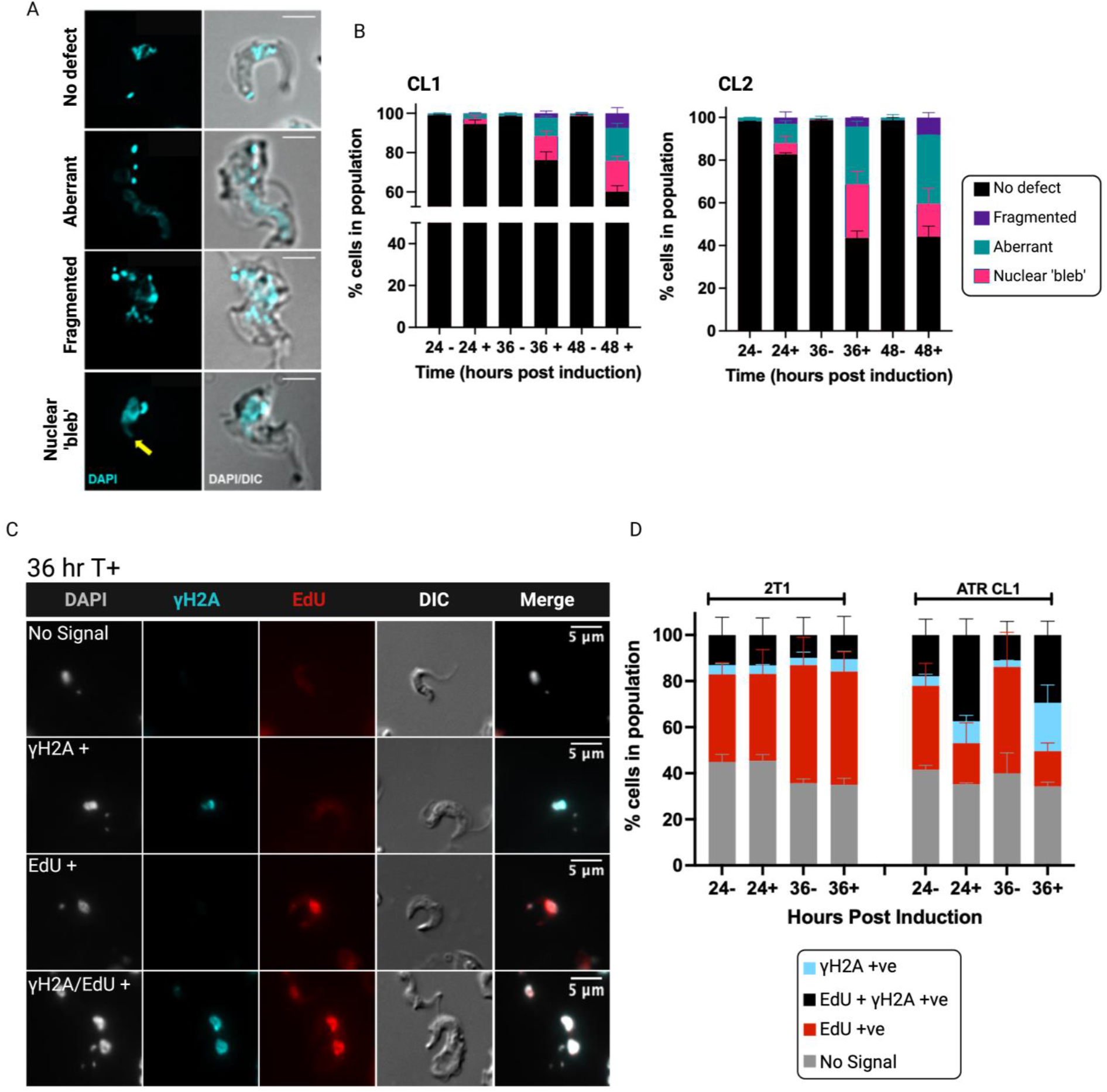
ATR loss affects nuclear division in *T. brucei* BSF cells but does not prevent DNA replication in the presence of genotoxic stress. (A) Representative examples of four categories of nuclei observed 36 hrs after TbATR RNAi; nuclear and kinetoplast DNA are stained with DAPI (cyan) and the cell body visualised by DIC. Scale bars = 5 μm. (B) The nuclei of BSF cells were examined after 24, 36 and 48 hrs growth in the presence (+) or absence (-) of TbATR RNAi induction by DAPI staining, and the number of cells in each of the four categories is expressed as a percentage of the total number of cells counted. Error bars = ± SEM; n=3 independent experiments for CL1, and n=2 for CL2 (>200 cells were counted/experiment). Significance was determined using a one-way ANOVA with Sidak’s multiple comparisons test (see Table S1) (C) Representative images of cells 36 hrs after RNAi induction (T+) and with EdU (red) and yH2A (cyan) labelling; DNA is stained with DAPI (grey). Scale bars = 5 µm. (D) Cells were scored for the presence or absence of EdU incorporation yH2A signal after growth for 24 and 36 hrs growth with (+) and without induction of RNAi in TbATR CL1 cells and 2T1 (control) parental cells. Values are expressed as a percentage of the total number of cells counted. Error bars = ± SEM; n=2 independent experiments (>100 cells counted/experiment).

### TbATR depleted cells continue to synthesise DNA

In addition to accumulating morphologically aberrant nuclei, TbATR depleted BSF cells also display increased levels of DNA damage markers, including increased nuclear yH2A levels and increased RPA and RAD51 foci^34^. To understand how abnormal nuclei formation after TbATR loss might relate to DNA replication, and to ask if damage accumulation prevents DNA replication, we tested if TbATR RNAi induced cells can synthesise nuclear DNA. To do this, we induced TbATR RNAi by addition of tetracycline and then pulsed them for 4 hrs with five-ethynl-2’-deoxyuridine (EdU; a thymidine analogue) prior to the desired timepoint. Cells were harvested after 24 and 36 hrs growth, including without induction of TbATR RNAi, and samples co-stained for EdU (using click-it chemistry) and yH2A (by indirect immunofluorescence^62^). EdU and yH2A signals were visualised by fluorescent microscopy and cells scored into four categories: no signal; with only EdU signal (EdU +ve); with only yH2A signal (yH2A +ve); and with both EdU and yH2A signal (EdU and yH2A +ve). Fig.1C shows representative images from CL1 collected 36 hrs after RNAi. Both in the parental RNAi cell line grown in tetracycline (2T1^46^, which does not target any gene by RNAi; Fig.S2A), and in the uninduced TbATR cells, most (∼35-40%; Fig.1D) of the population incorporated EdU and did not display yH2A signal, while ∼15-20% of cells displayed both signals, with just ∼5% of cells being only yH2A positive. In contrast, after TbATR RNAi, most cells (∼30-40%) co-stained with EdU and yH2A, suggesting DNA synthesis was proceeding despite increased genome damage (representative fields of view are shown in Fig. S2B). However, at both 24 and 36 hrs after RNAi increased numbers of cells displaying only yH2A signal were detected, perhaps indicating some cells accumulate sufficient levels of damage after TbATR loss to block continued DNA synthesis. Notably, the observation in both RNAi uninduced cells and parental 2T1 cells of co-staining for EdU and yH2A (∼20%) is consistent with prior studies suggesting the lack of a stringent checkpoint in *T. brucei* BSF cells to detect aberrant DNA during replication^48^. TbATR may then act only to signal the presence and limit the transmission of severely aberrant DNA.

### TbATR deficient cells are defective in chromosome segregation

To test further the impact of TbATR loss on chromosome segregation, we next examined telomere localisation using Telomere Fluorescence *In Situ* Hybridisation (Telo-FISH). Cells were collected after at 24 and 36 hrs growth with or without TbATR RNAi induction and telomere signal examined using multiple images acquired with deconvolved fluorescent microscopy. In addition to 11 megabase chromosomes, *T. brucei* harbours numerous mini- and intermediate chromosomes^44,63^ (containing ∼200 telomeres), and therefore the majority of signal detected in Telo-FISH is likely from the smaller chromosomes. In uninduced cells we observed cell cycle-dependent changes in telomere signal consistent with earlier studies^64,65^ (Fig.2A), with fluorescence detected around the nuclear periphery in G1, moving towards the centre of the nucleus in S-phase, and localising along the centre of the cell in metaphase. As mitosis progressed, and nuclear segregation occurred, the telomere signal re-localised to the nuclear periphery. Representative images of RNAi induced cells, at 36hrs, are shown in Fig.2B. To test for defective genome segregation, we quantified the number of distinct telomere foci present per nuclei (Fig.2C). In the uninduced population, the majority of cells harboured between 3-6 distinct telomere foci per nucleus. 36 hrs after RNAi this distribution of foci changed, with most cells displaying 4 or fewer foci. Strikingly, ∼30% of the induced population harboured no discernible telomere signal (increased from ∼5% of uninduced cells), suggesting a greater number of cells that failed to inherit nuclear DNA content. Analysis of the Telo-FISH images added to these data, as cells undergoing nuclear segregation were detected that did not appear to contain telomeric signal at the expected locations in the nucleus (Fig.1C; super resolution images are shown in Fig.S3). Furthermore, and consistent with DAPI staining (Fig.1), Telo-FISH signal could be detected that localised to small, DAPI stained structures, suggesting fragmentation of the nucleus. Taken together, these data indicate that TbATR loss not only impairs nuclear structure but also undermines accurate chromosome segregation.

**Figure 2.**
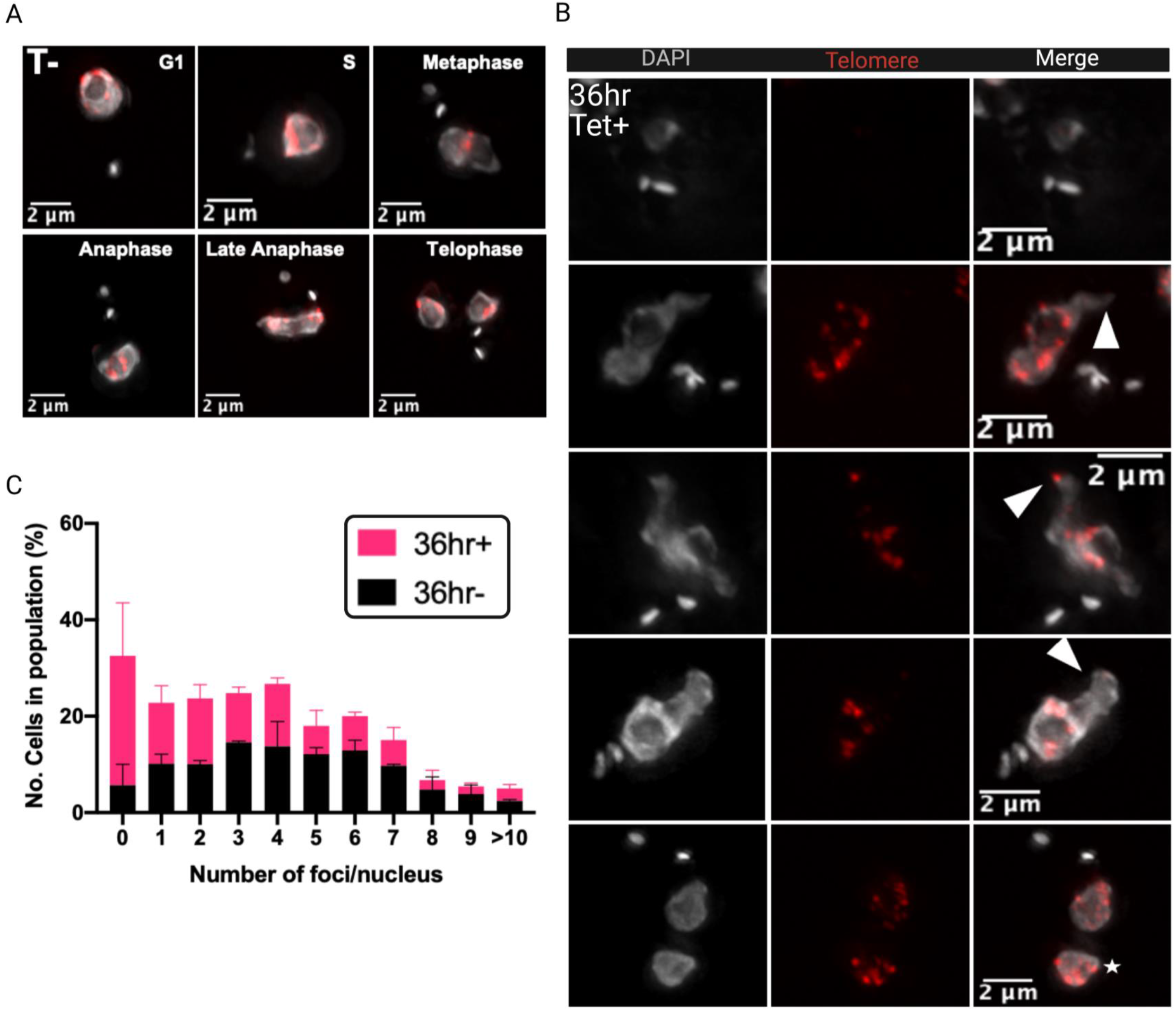
Depletion of ATR in *T. brucei* BSF cells disrupts chromosome segregation. (A) Representative images of telomere localisation in control, uninduced cells (T-; 36 hrs) throughout the cell cycle. The nuclei were stained with DAPI (grey) and the telomeres localised using a telomere probe conjugated with FITC (Telo-FISH; red). Scale bars = 2 µm. (B) Representative images of cells 36 hrs after induction of RNAi against TbATR (Tet+). White arrows indicate uneven segregation of telomeric foci; white star indicates a cell in which one nucleus harbours a greater number of telomeric foci signal than the other. Scale bars = 2 µm. (C) Cells were collected after 36 hrs growth with (+) and without (-) TbATR RNAi induction and the number of distinct Telo-FISH foci in each nucleus counted. Error bars = ± SEM; n=2 independent experiments (>100 cells counted/experiment).

### TbATR loss causes widespread changes in transcription

We previously reported that TbATR depletion rapidly and significantly undermines the control of RNA Pol I gene expression in BSF cells, most notably due to increased transcription of normally silent variant surface glycoprotein (VSG) genes required for immune evasion by antigenic variation^34^. A small cohort of RNA Pol II transcribed genes also displayed altered RNA levels 24 hrs after TbATR RNAi, though none showed the same level of change. These data hint at wider effects of TbATR loss on *T. brucei* gene expression. To test this, and to ask if changes in expression of genes with functions in nuclear maintenance and cell cycle progression were seen that may reveal TbATR signalling of damage, we performed differential RNA-seq, comparing gene-specific RNA levels in BSF cells grown for 36 hrs with and without RNAi induction (Fig. 3, Tables S3, S4; induced and uninduced samples were collected in triplicate, from separate experiments).

**Figure 3.**
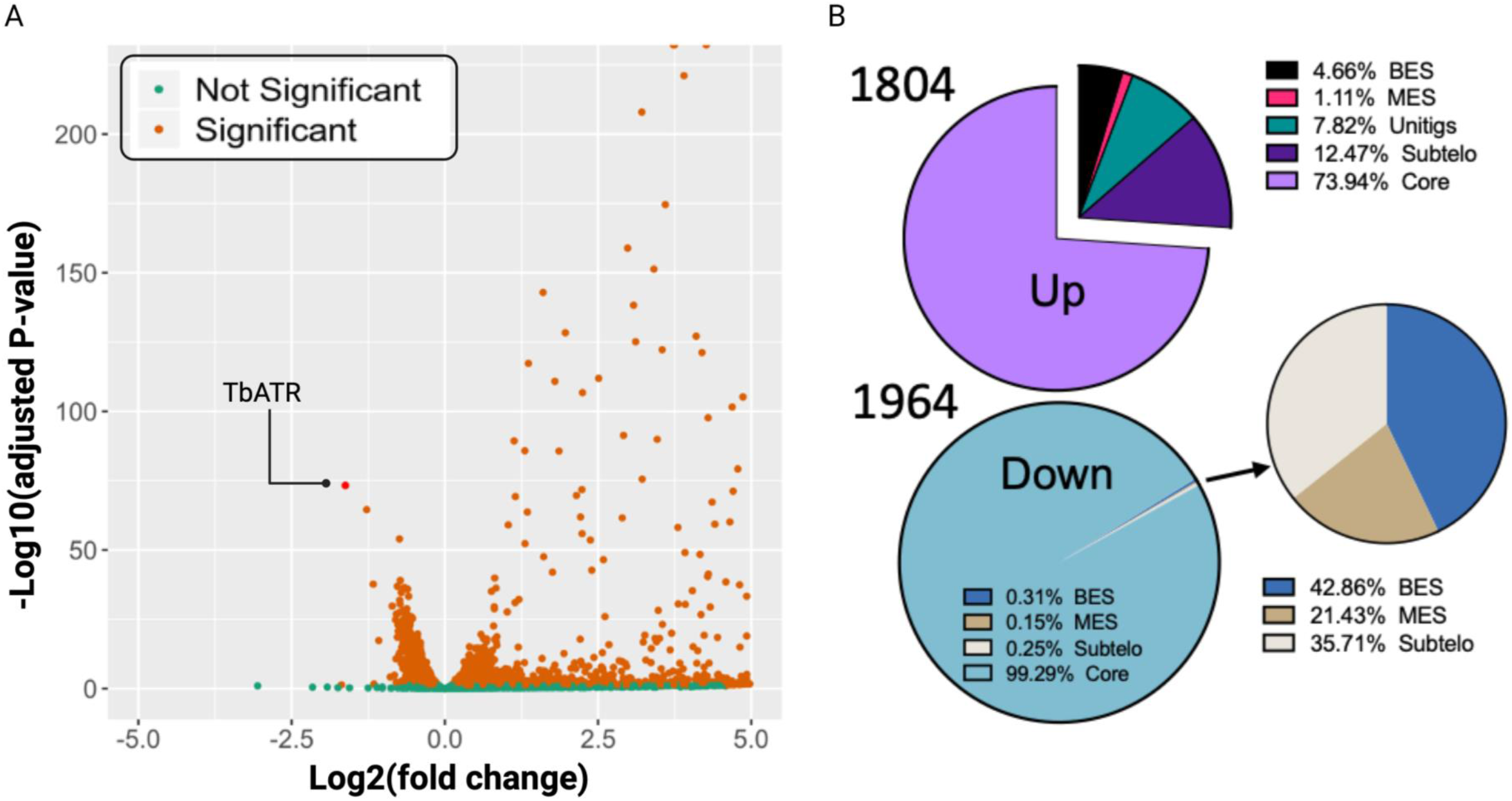
ATR RNAi leads to widespread changes in *T. brucei* BSF gene expression. (A) A Volcano plot showing differentially expressed transcripts 36 hrs after RNAi relative to uninduced controls. Log10 adjusted p-values for each gene are plotted against the log2 transformed fold-change; data are averages from three biological replicates and genes showing a significant change in abundance are in orange (non-significant in green); TbATR is indicated. (B) Pie charts summarising differentially expressed transcripts (up, increased; down, decreased) 36 hrs after RNAi; the number of genes in each category is expressed as a percentage of total gene number, and genes were categorised based on their genomic location (core genome; BES; MES, subtelomere and unmapped unitigs). The smaller, lower piechart shows the genomic distribution of significantly differentially expressed non-core genes with reduced RNA levels after RNAi.

In total, 3768 significantly differentially expressed transcripts were identified 36 hrs after TbATR RNAi (significance determined by an adjusted p-value < 0.05; Fig.3), meaning 42% of predicted protein-coding genes displayed altered expression. A small majority of genes (1964, 52%; Table S3) displayed significant reductions in RNA levels, though the extent of change was greater in the minority (48%, 1804; Table S4) of genes whose RNA levels significantly increased. Consistent with changes in RNA Pol I expression 24 hrs after RNAi^34^, the majority of differentially expressed RNA Pol I transcribed genes whose RNA levels increased were located in silent BES (∼80%: 84 of the 104 significantly differentially expressed RNA Pol I transcribed genes identified; Table S3). Moreover, nearly all down-regulated RNA Pol I genes were located in the single actively transcribed VSG bloodstream expression site (BES1; Table S4; with the exception of one expression site associated gene (ESAG) from a silent BES (BES13)). The remaining differentially expressed RNA Pol I transcribed genes were located within metacyclic expression sites (MES), which provide the metacyclic life cycle stage with a protective VSG surface coat. Thus, loss of telomeric VSG ES control continues upon prolonged TbATR depletion^34^. The striking difference revealed by this differential RNA-seq of TbATR RNAi relative to 24 hrs, was that after 36 hrs RNAi the majority of differential expression was attributed to genes located within the core of the genome (>99% of those down-regulated, and ∼74% of up-regulated; Fig.3), indicating very widespread disruption of RNA Pol II transcription. Reinforcing the potentially global gene expression alteration due to loss of TbATR, GO term enrichment analysis revealed a wide range of predicted functions, with no clear evidence that DNA repair, replication or signalling genes (which may have been predicted to change due to loss of a DNA damage signalling kinase) were a particular focus of change (Figure S4).

### Depletion of TbATR is associated with damage accumulation in intergenic and subtelomeric regions of the genome

To understand why TbATR depletion should have such widespread effects on gene expression, we next examined the pattern of DNA damage in the genome after RNAi. Previously, we documented that TbATR RNAi for 24 hrs is associated with the accumulation of yH2A signal^62^ within the VSG BES. To investigate the distribution of yH2A following 36 hrs TbATR RNAi, we mapped yH2A ChIP-seq reads across all 11 *T. brucei* megabase chromosomes (Fig.4). By comparing yH2A enrichment at 24 and 36 hrs after RNAi induction relative to uninduced, it became apparent that whereas there was little accumulation of the phosphorylated histone within RNA Pol II multigene transcription units after 24 hrs RNAi, pronounced accumulation was seen between CDS after 36 hrs (Fig.4A). Metaplots comparing all RNA Pol II transcribed protein-coding genes (Fig.4B) confirmed this prediction, with clear peaks of yH2A ChIP-seq reads seen upstream and downstream of CDS after 36 hrs of TbATR RNAi, and no such peaks after 24 hrs. Such yH2A enrichment after TbATR loss was not limited to protein-coding genes but was also seen at rRNA loci and may, in fact, arise earlier at these RNA Pol I units (Supplementary File 1). In addition, accumulation of yH2A from 24-36 hrs after TbATR RNAi was also seen at the initiation and termination sites of the RNA Pol II transcription units, though the level of enrichment was greater at transcription start sites (Fig.S5A).

**Figure 4.**
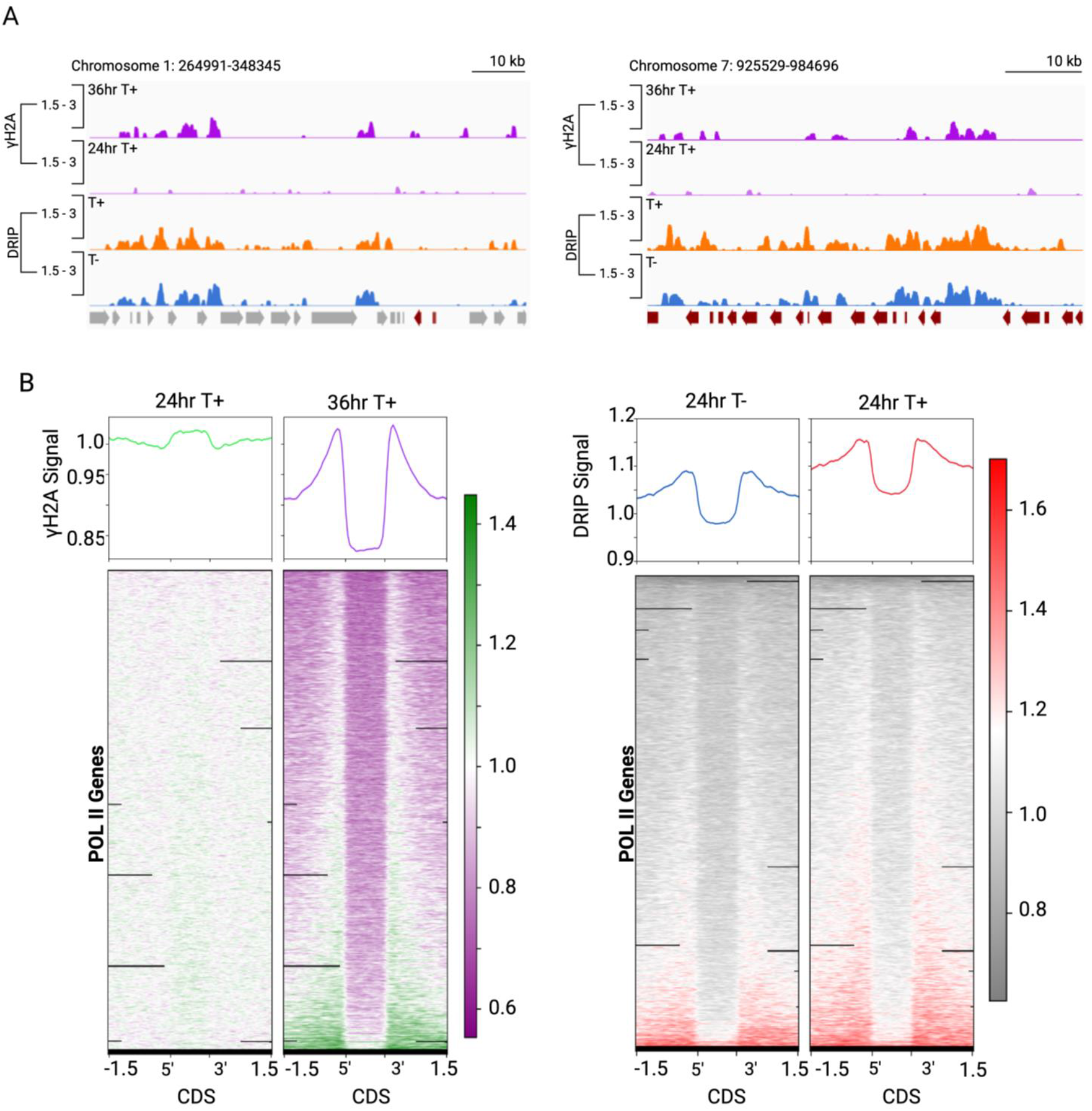
R-loops and DNA damage colocalise at intergenic genome regions within *T. brucei* multigene RNA Polymerase II transcription units after ATR RNAi. (A) Representative plots showing yH2A ChIPseq and S9.6 DRIPseq signal across RNA Polymerase (Pol) II transcribed regions of chromosomes 1 and 7. yH2A ChIP-seq signal enrichment (y axis) is shown as a ratio of reads in RNAi induced (T+) samples relative to uninduced (T-; each normalised to cognate input sample) after growth for 24 and 36 hrs; DRIPseq signal is shown as level of enrichment in DRIP relative to input after 24 hrs growth with (T+) and without (T-) RNAi; data was visualised using Gviz. (B) Metaplots and corresponding heatmaps: left panel, normalised yH2A signal 24 and 36 hrs post TbATR RNAi plotted across all predicted RNA Pol II transcribed CDSs; right panel, normalised DRIPseq after 24 hrs growth with (+) and without (-) TbATR RNAi; in both panels each CDS is scaled to 1.0 Kb and data is plotted +/- 1.5 Kb.

In previous studies^52,66^ we showed inter-CDS regions and transcription start sites are sites of R-loop accumulation, and the RNA-DNA hybrids in these locations are acted upon by RNase H enzymes. Thus, to ask if the loss of TbATR and the corresponding yH2A accumulation could be attributed to R-loops, we used DRIP-seq^52^ with S9.6 antibody to immunoprecipitate and map the locations of RNA-DNA hybrids before and after induction of TbATR RNAi. Reads in both the 24 hr RNAi-induced and uninduced DRIP samples were mapped as a fold change in abundance relative to their corresponding input samples (Fig.4); for reasons that remain unclear, DRIP was unsuccessful 36 hrs after TbATR RNAi.

Plotting DRIP-seq signal across the chromosome cores showed the accumulation of R-loops in the absence of RNAi induction that matched previous observations in WT cells^52^, with signal enrichment at inter-CDS regions within multigene transcription units (Fig.4A), as well as at regions of transcription initiation (Fig.S5B). Strikingly, the locations of yH2A ChIP-seq enrichment 36 hrs after TbATR RNAi closely matched the DRIP-seq (Fig.4A) with metaplots showing both signals as very similar peaks upstream and downstream of CDS (Fig.4B). Thus, there was genome-wide correlation between DNA damage localisation after loss of TbATR and R-loop distribution in the multigene transcription units of the genome. Such correlation was less clear at transcription initiation sites, where yH2A ChIP-seq signal appeared more broadly distributed than the more localised DRIP-seq accumulation (Fig.S5). Comparing DRIP-seq enrichment in the RNAi induced and uninduced cells (Fig.4B) suggested a modest increase in R-loops around the CDS after TbATR was depleted by RNAi for 24 hrs, with a similar effect seen at sites of transcription initiation (Fig.S5). Though a distinct effect on DRIP-seq enrichment after TbATR loss at subtelomeric and centromeric loci (see below) further suggests that loss of the kinase alters R-loop abundance and distribution, the true extent of such changes may be obscured by data normalisation during mapping.

### Correlation between R-loops and DNA damage within BES after TbATR depletion

We previously reported that loss of TbATR leads to yH2A accumulation in the VSG BES, causing elevated levels of VSG expression switching^34^. In addition, we have shown that loss of RNase H1^67^ or RNase H2A^66^ causes accumulation of R-loops in the VSG BES, also leading to yH2A accumulation and increased VSG switching. To ask if the VSG BES DNA damage after TbATR loss correlates with R-loops, as was seen in the chromosome core, DRIP-seq was mapped to the active and silent loci, with and without TbATR RNAi using MapQ filtering to limit cross-mapping to these homologous sites (Fig.5, Fig.S6). As expected, accumulation of yH2A was seen across the active and silent BES, with greater signal around the VSG (associated with the 70 bp repeats and telomeres) 36 hrs after RNAi. R-loops were detected in both active and silent BES even in the absence of TbATR RNAi induction, which contrasts with WT cells but is comparable with uninduced RNase H2A RNAi cells^66^, perhaps indicating leakiness in the TbATR RNAi. These data suggest that a modest loss of TbATR results in R-loop accumulation, as seen in the chromosome core after RNAi induction. However, after TbATR RNAi, whereas R-loop levels appeared unchanged across the ESAG-encoding sequence of the active and silent BES, reduced R-loop enrichment was detected across the 70 bp repeats and downstream of the VSG gene, a localised effect that was consistent in all BES (metaplot, Fig.5B) and coincided with increased yH2A signal at the same BES sequences (Fig.5A, S6). Thus, though some R-loops in the RNA Pol I transcribed VSG BES appear to be recognised by TbATR and lead to DNA damage after loss of the kinase without substantial change in abundance of the RNA-DNA hybrids, TbATR loss appears to have a more pronounced effect on damage levels and to result in distinct R-loop processing or stability on sequences around the telomere-proximal VSG.

**Figure 5.**
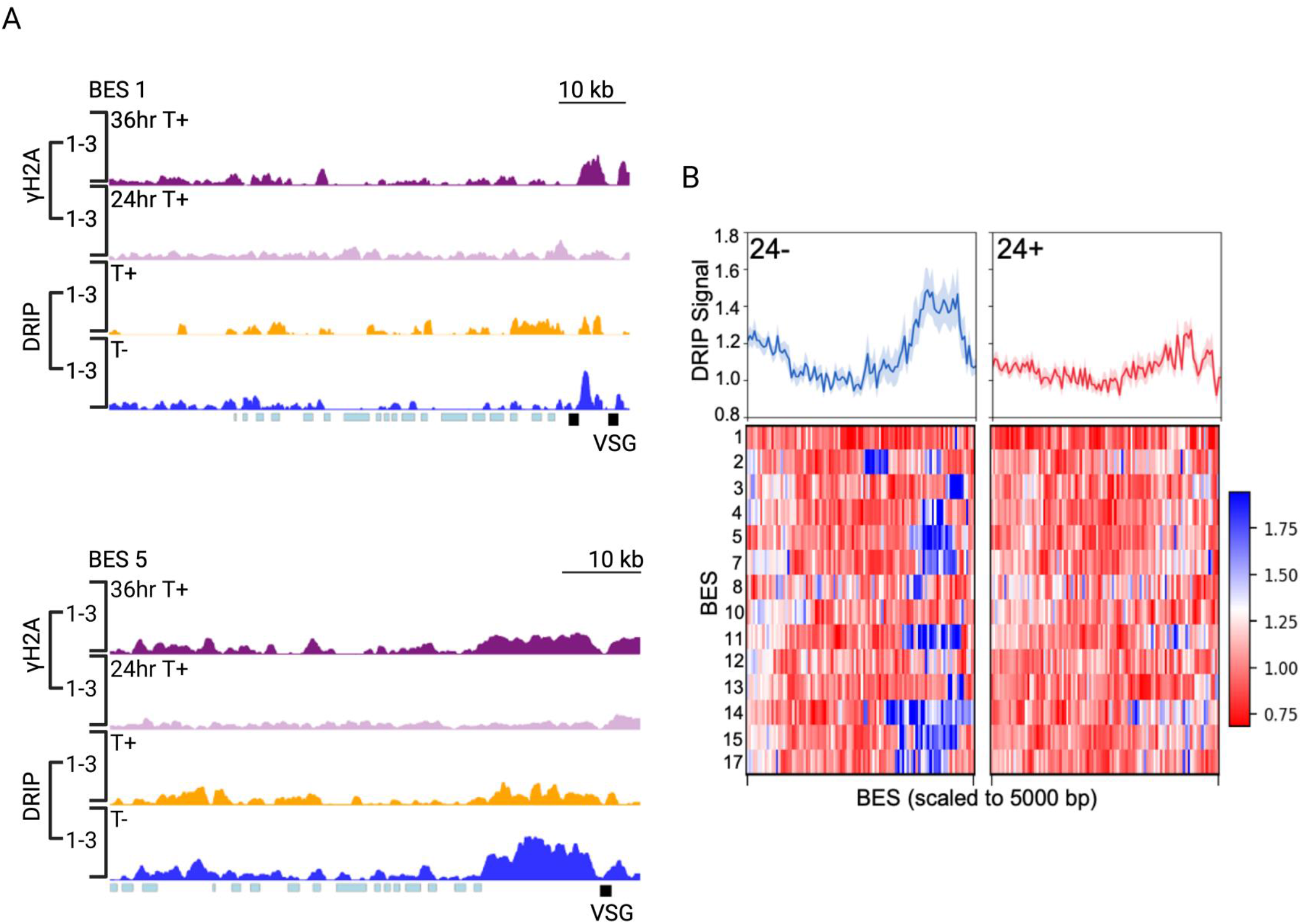
Accumulation of DNA damage after ATR RNAi in *T. brucei* VSG expression sites (BES) correlates with reduced R-loop detection. (A) Normalised yH2A and DRIPseq signal plotted across the whole of the active BES (BES1; containing VSG2) and one silent BES (BES5). Data are shown and plotted as in Fig.4; VSG genes are shown as dark blue boxes and upstream expression site associated genes as light blue boxes. (B) Metaplots and heatmaps show the distribution of DRIPseq signal after 24 hrs growth with (24+) and without (24-) TbATR RNAi induction across all BES; BES number is shown to the left of the heatmap, and each BES is scaled to the same size. Mean DRIPseq signal (DRIP enrichment normalised to input) is represented as a solid line and standard error is shown as a shaded area.

### TbATR acts on centromeric R-loops in *T. brucei*

R-loops are also readily detected within *T. brucei* centromeres^52^, though how and why RNA-DNA hybrids form at these loci is unknown. Given that TbATR depletion led to impaired chromosome segregation (Fig.2), we asked if TbATR RNAi altered R-loop and yH2A abundance across all annotated centromeres in the *T. brucei* genome, as defined by the localisation of kinetochore component TbKKT2^44^ (Fig.6, S7). In keeping with prior observations, we detected DRIP-seq read accumulation across all but one centromere in the absence of TbATR RNAi induction. Furthermore, there was no evidence for enrichment of yH2A ChIP-seq reads within centromeres 24 hrs after TbATR RNAi. However, 36 hrs after RNAi induction we observed accumulation of centromeric yH2A ChIP signal in 10 of the 14 predicted centromeres (Fig.6B), with the extent of enrichment broadly correlating with R-loop levels in the uninduced cells (as predicted from DRIP-seq). In contrast, a modest reduction in centromeric DRIP-seq enrichment was seen after TbATR depletion by RNAi for 24 hrs (Fig.6C). As discussed above, we note that caution should be taken in assessing these changes in levels between samples, given data normalisation. Nonetheless, these data support a parallel effect of TbATR RNAi at the centromeres to that observed in telomere-proximal regions of the VSG BES: loss of the kinase leads to damage after 36 hrs, which appears to follow a modest loss of R-loops. Hence, TbATR directs R-loop homeostasis at *T. brucei* centromeres and absence of the kinase leads to DNA damage.

**Figure 6.**
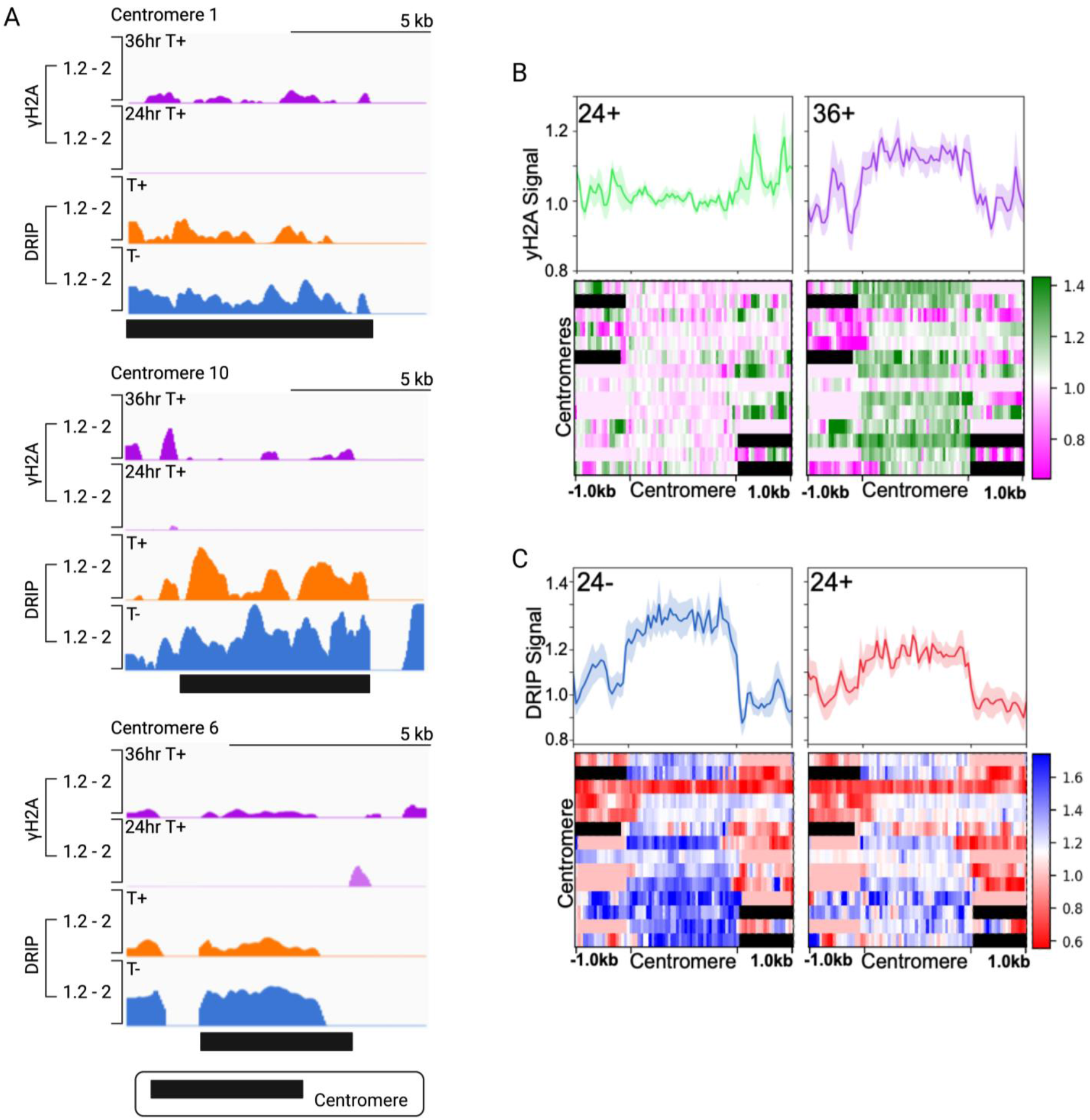
Accumulation of DNA damage after ATR RNAi in *T. brucei* centromeres correlates with reduced R-loop detection. (A) Normalised yH2A and DRIPseq signal plotted across centromeric proximal regions of chromosomes 1, 10 and 6; data are shown and plotted as in Fig.4 and centromere location is marked by a black box. (B) Metaplots and corresponding heatmaps of normalised yH2A signal 24 and 36 hrs post TbATR RNAi plotted across all centromeres; each centromere is scaled to 2.0 kb and data is plotted with +/- 1.5 Kb of surrounding sequence. Mean signal is represented as a solid line and standard error is shown as a shaded area. (C) Similarly represented metaplots and corresponding heatmaps of DRIPseq after 24 hrs growth with (+) and without (-) TbATR RNAi (DRIP enrichment normalised to input).

### Loss of TbATR leads to impaired spindle formation during mitosis

Recent work in human cells implicates a role for centromeric R-loops in directing an ATR-mediated pathway during mitosis to enable faithful chromosome segregation^26^. Thus, to test if altered centromeric R-loop levels and increased DNA damage after TbATR loss might explain impaired chromosome segregation in *T. brucei*, we examined TbATR RNAi induced and uninduced cells for the presence or absence of mitotic spindles. Cells were collected after at 24 and 36 hrs growth (Fig.7 and Fig.S8) and mitotic spindles examined by indirect immunofluorescence using anti-tubulin KMX-1 antiserum. Nuclear and kinetoplast DNA were visualised by DAPI staining. Representative images of cells after a 36 hrs growth and containing nuclear spindles are shown in Fig.7A,B (uninduced and induced, respectively; an antibody negative control is shown in Fig.S8A) and are quantified in Fig. 7C. In uninduced cells, ∼12 % of the population possessed a mitotic spindle, in keeping with previous findings^68^ and correlating with numbers of 1N2K cells in the population^34^. Following depletion of TbATR a significant reduction in the number of cells possessing a recognisable mitotic spindle (∼5%) was seen. This loss appears especially striking, because TbATR RNAi causes accumulation of 1N2K cells, which are in the cell cycle stage in which mitotic spindles are readily detected^34^. Furthermore, though spindle loss across the population could be explained by an increased number of cells with aberrant nuclear configurations, we also observed a significant reduction in the number of cells with recognisable mitotic spindles (Fig.S8B) 24 hrs post RNAi induction, when 1N2K cells accumulate but very few cells with aberrant nuclear configurations are seen^34^. Thus, our data suggest TbATR RNAi leads to defective mitotic spindle assembly in BSF *T. brucei*, which may then lead to chromosome and nuclear segregation defects.

**Figure 7.**
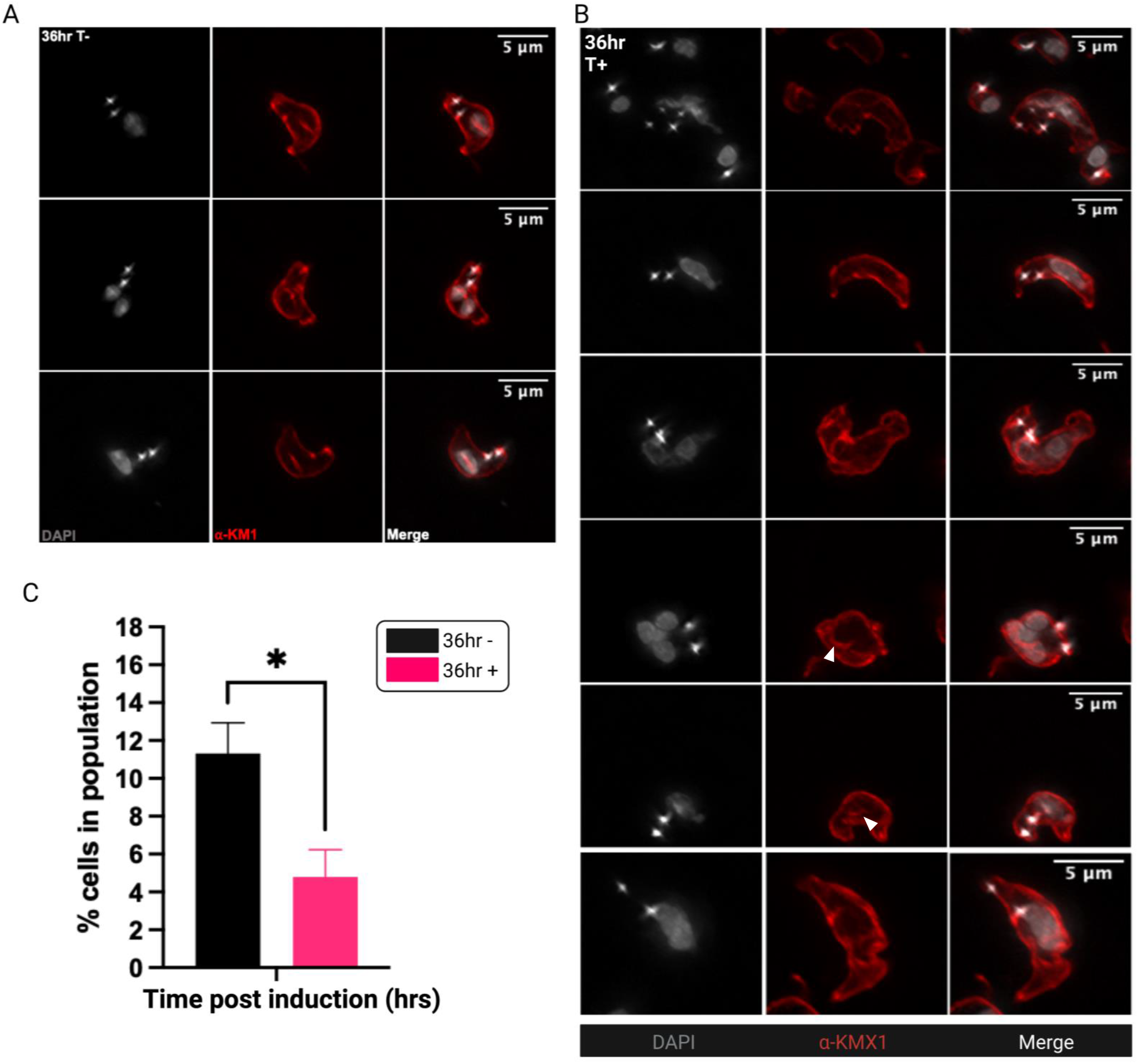
ATR RNAi in *T. brucei* BSF cells leads to impaired mitotic spindle formation. (A) Representative images of cells after 36 hrs growth without (T-) TbATR RNAi and after mitotic spindle labelling using tubulin antiserum anti-KMX-1 (red). Nuclear and kinetoplast DNA were stained with DAPI (grey). Scale bars = 5 µm. (B) Representative images of cells after 36 hrs growth with induction of TbATR RNAi (T+). (C) Cells were collected after 36 hrs growth with (36 hr +) and without (36 hr-) TbATR RNAi induction and mitotic spindle detected via indirect immunofluorescence (as above); the number cells in the population containing a recognisable mitotic spindle were counted and are shown as a percentage of the total number counted. Error bars = ± SEM; n=3 independent experiments (>100 cells counted/experiment). (*) p = 0.0393, unpaired t-test (two-tailed).

### TbATR localises as a thread between segregating nuclei during mitosis

The data above suggest that TbATR provides hitherto undetected functions in chromosome segregation in BSF *T. brucei*. To test this further, and to ask if we could detect a role during mitosis, we sought to localise the protein in the cell. To do so we added a 12myc epitope tag to the N-terminus of one *TbATR* allele, and confirmed integration and expression by PCR and western blotting (Fig.S9A-C). Deletion of the untagged allele and analysis of growth and sensitivity to MMS indicated that the 12myc-TbATR tagged variant is functional (Fig.S9D). We next localised TbATR by indirect immunofluorescence using anti-myc antiserum. Nuclear and kinetoplastid DNA were visualised by DAPI staining. Anti-myc signal was present in the nuclei of cells across all stages of the cell cycle (Fig. 8, Fig. S9E-F). Additionally, in mitotic cells, we could detect myc signal spanning the area between the two segregating nuclei (Fig.8A, an antibody negative control is shown in Fig. 8B), which was clearly visible after structured-illumination super-resolution imaging (Fig.8C). Thus, while TbATR is a nuclear protein, it also found in a thread-like bridge between segregating nuclei during mitosis of BSF *T. brucei* cells (Fig 8 and Fig. S9G). Such a localisation could be consistent with a function for TbATR during chromosome segregation.

**Figure 8.**
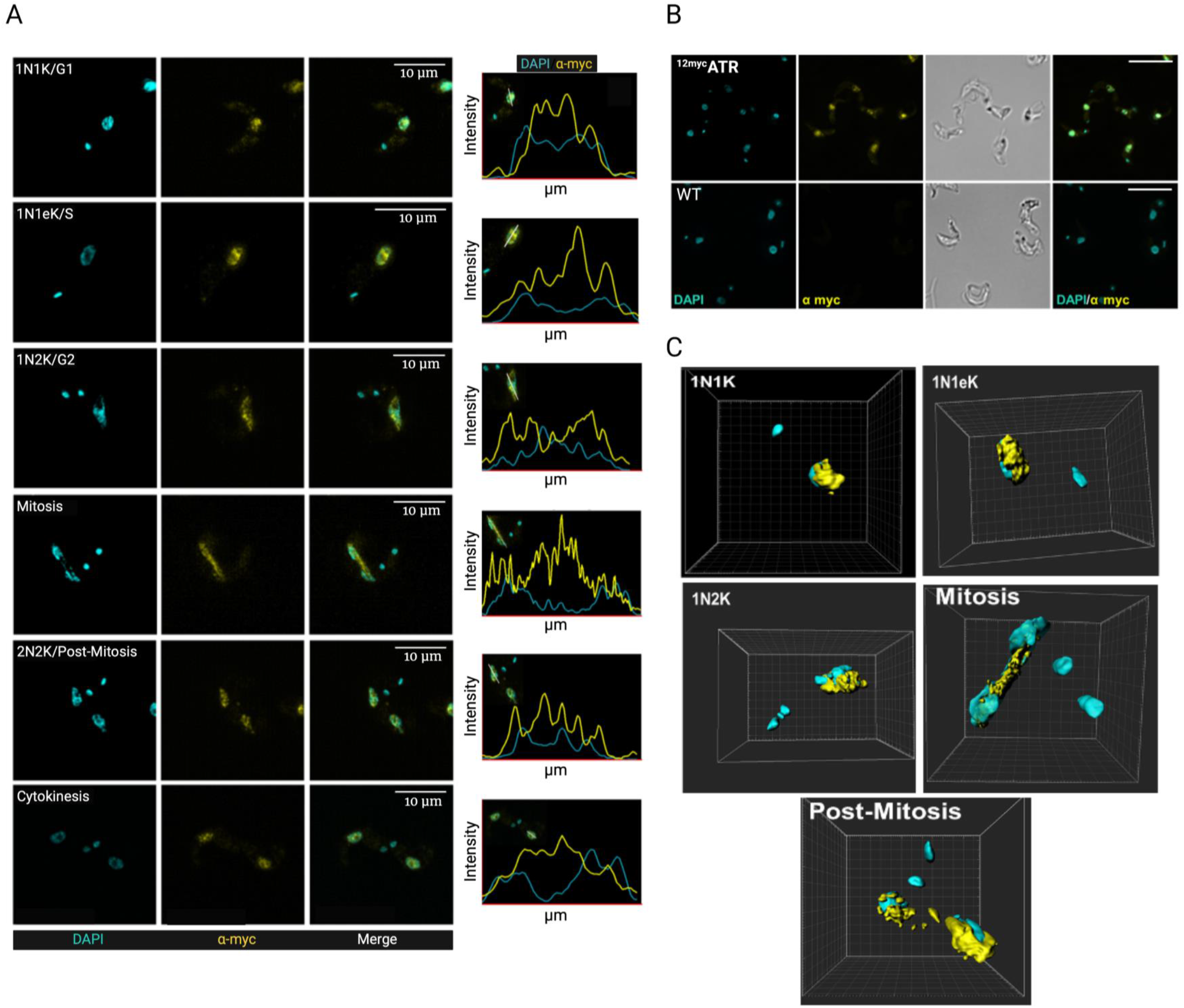
T. brucei ATR localises in a thread-like manner between segregating nuclei during mitosis. (A) Representative images of the sub-cellular localisation of myc signal in cells expressing TbATR N-terminally tagged with 12 myc (^12myc^ATR+/- cells); cells at different stages of the cell cycle are shown; scale bar = 10 µm. Images were captured on an Elyra super resolution microscope (Zeiss). Nuclear (n) and kinetoplast (k) DNA are shown in cyan, and myc signal in yellow. Graphs to the right show fluorescence intensity (arbitrary units) of the two signals across a region of the cells (µm) marked by the white line. (B) A comparison of the sub-cellular localisation of the myc signal in ^12myc^ATR+/- and WT (untagged, control) cells. Images were captured on a DeltaVision microscope. Scale bar = 20 µm. (C) 3D rendered images of ^12myc^ATR+/- cells in different cell cycle stages generated from super resolution stacks captured on an Elyra super resolution microscope (Zeiss). Images were generated in IMARIS: nuclear and kinetoplast DNA is in cyan, and myc signal in yellow.

## Discussion

Conservation of the ATR kinase across most eukaryotes highlights its fundamental role in genome maintenance and transmission. Often cited as the ‘master regulator’ of the DDR, ATR acts to survey and signal replication-derived lesions by responding to the accumulation of ssDNA^16^. More recently, wider roles for ATR have been recognised, including maintaining the integrity of the nuclear envelope^27,29,30^, nucleolus^28^ and centrosome^31^. In the kinetoplastid parasite *T. brucei*, TbATR acts in cell cycle progression and DNA double-strand break repair^37^, and in mammal-infective BSF cells the kinase acts in monoallelic VSG expression and VSG recombination during host immune evasion by antigenic variation^34^. However, whether such VSG-related functions could explain why loss of TbATR is essential in mammal-infective *T. brucei*, but not in tsetse-infective cells, is questionable. Here, we report widespread R-loop associated roles for TbATR that appear critical for chromosome segregation and gene expression, revealing both conserved and lineage-specific essential functions of ATR.

Consistent with TbATR having wide roles in *T. brucei* biology - akin to the emerging data on diverse ATR functions in other eukaryotes - analysis of various cellular phenotypes reveals more pleotrophic effects after its loss than the impaired cell cycle progression, increased DNA damage and altered VSG expression control documented previously^34,35,37^. First, TbATR depleted cells display numerous changes in their nuclear ultrastructure. Whether or not these effects reflect a conserved role in maintaining nucleus integrity^30^, and whether such a surveillance function extends to monitoring the nucleolus^28^ or is executed via nuclear F-actin^69^, is unclear, although loss of the kinase appears to alter localisation of at least one subnuclear structure, the VSG expression site body^34^. Second, we document that *T. brucei* cells depleted of TbATR fail to segregate their nuclear genome correctly, including the emergence of unevenly divided nuclear chromatin and, more severely, cells with nuclei lacking telomeric signal detectable by Telo-FISH. Unusually, we also detect aberrant BSF *T. brucei* cells known as zoids ^34,35,70^, which stain only for mitochondrial DNA, following TbATR depletion, indicating wholesale loss off the nucleus after mitosis. These phenotypes can be partially explained by the observation that loss of TbATR does not prevent nuclear DNA replication, including in the presence of increased levels of yH2A, which may be consistent with TbATR-dependent checkpoint signalling of genome damage only partially preventing progression of mitosis in BSF *T. brucei*^48^. Third, we show that loss of TbATR alters R-loop accumulation at centromeres and impairs spindle formation during mitosis, suggesting a mechanistic connection between RNA-DNA hybrids and TbATR activity during chromosome segregation that is distinct from damage signalling, and may be consistent with the demonstration that the kinase localises between nuclei as they separate during mitosis. Finally, we demonstrate that TbATR RNAi has very widespread effects on *T. brucei* gene expression: the earliest and most pronounced changes are seen at telomeric RNA Pol I-transcribed VSG BES, but by 36 hrs nearly half of RNA Pol II transcribed genes also display significantly altered RNA levels. Such widespread perturbation in gene expression seems unlikely to merely reflect TbATR signalling of DNA repair activities. Instead, our data indicate that loss of TbATR affects multigene transcription, since RNAi of the kinase results in increased yH2A signal at inter-CDS regions across the genome, colocalising with previously mapped sites of R-loop accumulation^52^.

Previous work has clearly implicated ATR in the response of both *T. brucei* and *Leishmania* to DNA damage: loss or inhibition of the kinase sensitises the parasites to lesions resulting from alkylation^45^, irradiation^37^ and oxidative stress^71^; in addition, TbATR RNAi in *T. brucei* results in focal accumulation of RPA and RAD51, and increased levels of yH2A, which have been mapped to the VSG expression sites^34^. Nonetheless, the action of ATR in kinetoplastid DNA replication has not been as clearly described, and its role in preventing the segregation of damaged chromosomes may not be absolute, at least in *T. brucei*. Two key roles of ATR in other eukaryotes are modulating S/G2^4^ and G2/M^72^ checkpoints to allow repair of genome damage prior to mitosis, and monitoring DNA replication through responding to replication fork-blocking lesions and activation of backup origins^16^. In BSF *T. brucei*, alkylation damage by methyl methanesulphonate does not elicit a clear cell cycle checkpoint, though loss of a number of kinases, including TbATR, causes increased damage sensitivity^45^. Even more strikingly, DNA double-strand breaks induced at either telomeric or chromosome-central loci have been shown to persist through mitosis without repair, as monitored by the presence of persistent RPA foci^48^. Thus, S/G2 or G2/M checkpoints appear incomplete, at best. Why and how *T. brucei* has abrogated these checkpoints is unclear. One proposal is that damage does not always prevent cell division in *T. brucei* in order to promote genetic diversification and adaptation, such as during immune evasion^48^. If so, such checkpoint liberation may be maintained throughout the parasite’s life cycle, since whereas TbATR-directed S/G1 and intra-S checkpoints were detected after irradiation of PCF *T. brucei*, S/G2 and G2/M checkpoints were not described^37^. Moreover, these data suggest that TbATR directs DNA damage signalling in G1/S phases of the cell cycle, raising the question of how such activity is suppressed through G2-M phases. One problem in addressing these questions is incomplete understanding of the ATR pathway in *T. brucei*. For instance, no orthologue of CHK1 has been identified in any kinetoplastid, and though a kinase with similarity CHK2 (which acts downstream of ATM) has been proposed^38^, its functions are untested. Moreover, whereas 9-1-1 functions have been linked to genome variability and subtelomere replication in *L. major*^39,73^, no work has examined the role of the complex in *T. brucei*.

Whether or not the above divergence in TbATR activity during DNA repair extends to its roles in monitoring origin activity and replication fork progression during DNA replication in *T. brucei* is less clear. In both BSF and PCF *T. brucei*, TbATR RNAi or inhibition causes some accumulation of cells in S-phase^35,45^, indicative of an effect on DNA replication progression. However, no data so far have revealed if loss of TbATR affects the global programme of origin firing in *T. brucei*, as has been described in other eukaryotes^74–77^. We show here that EdU incorporation continues in BSF *T. brucei* cells after RNAi, indicating DNA replication does not halt, and previous work suggested hydroxyurea impairment of DNA replication increases activity from one origin^78^, though genome-wide analysis is lacking. Nonetheless, two observations suggest further work is needed. One observation is that the known signalling machinery for ATR modulation of origin activity is not obviously present in *T. brucei*, including CHK1 and CDC7 (DBF4-dependent) kinases^74,75^. Instead, in *T. brucei* a complex of CRK2 (Cdc2-related kinase 2) and CYC13 activates the replisome through phosphorylation of the MCM helicase^79^, and it is unclear if TbATR contributes to such activity. The second observation is genome-wide mapping of DNA replication dynamics in *L. major*, which indicates that hydroxyurea treatment does not detectably result in activation of DNA replication initiation at any site beyond the single predicted S-phase origin in each chromosome; in fact, the main effect of such replication stress is loss of extra-S phase subtelomeric DNA replication^73^. Thus, it is possible that kinetoplastids have evolved to copy their genomes without the safety net of back-up, dormant origins^80^.

An important role of ATR in tackling impaired replication fork progression is in dealing with conflicts between DNA replication and transcription^19,81,82^, which often result in R-loops. Such conflicts are especially prevalent in large genes, where transcription overlaps with DNA replication to a greater extent, causing DNA damage, fragility and R-loop accumulation being more common in such genes^83–85^. The organisation of the *T. brucei* genome into ∼200 multigene transcription units^86^, some containing hundreds of co-transcribed genes, might be interpreted as the parasite only encoding large genes, with the prediction of transcription-replication conflicts being prevalent and localised. Indeed, transcription is observed throughout the *T. brucei* cell cycle^87^, and mapping suggests DNA replication forks meeting multigene transcription head-on are slowed relative to those that are co-directional with transcription^88^. Despite these observations, previous mapping, with and without loss of RNase H1 or RNase H2, did not reveal accumulation of R-loops at predictable locations relative to origins^52^, and no such pattern was detected here after TbATR RNAi. Instead, we reveal that loss of TbATR causes pronounced accumulation of yH2A at potentially every site of R-loop enrichment within the multigene transcription units, corresponding with inter-CDS regions of pre-mRNA processing by trans-splicing and polyadenylation. Thus, the main transcription-associated role of TbATR appears to be monitoring R-loops that derive from regions of RNA Pol pausing, which may be the major impediments to replication fork progression in the *T. brucei* genome. Loss of TbATR in monitoring inter-CDS R-loops would explain the very widespread effects on gene expression after TbATR RNAi. Whether this role is mechanistically related to the recent description of ATR acting on R-loops during mammalian mRNA polyadenylation is unclear^89^, since polyadenylation in *T. brucei* mRNA maturation may be distinct, as it is intimately coupled with trans-splicing^90^. We also observed yH2A accumulation after TbATR RNAi at transcription initiation sites, which are the major locations of damage formation after loss of TbRNase H2^66^. Since loss of TbRNase H2 does not result in detectable inter-CDS yH2A signal, it is possible that R-loop processing and monitoring at sites of transcription initiation and processing are distinct. Nonetheless, interplay between R-loops and TbATR activity is not limited to RNA Pol II transcription, as we find the same pattern of damage accumulation after RNAi at VSG ESs and at rRNA loci, which are transcribed by RNA Pol I. A common and widespread mechanism of RNA Pol pausing causing impaired DNA replication fork movement, which is recognised and tackled by TbATR, might be predicted, but needs further analysis. Nevertheless, the effects we describe after TbATR RNAi provide further evidence that *T. brucei* immune evasion by VSG switching is elicited by a mechanism that links R-loop levels with transcription-replication conflict in the VSG expression sites^34,43,67,91^.

This work reveals a hitherto undetected role for TbATR in *T. brucei* chromosome segregation, which is also mediated through R-loops but appears distinct from the transcription-association function discussed above. We show that loss of TbATR causes accumulation of yH2A and changes in R-loop levels at centromeres and, in addition, impairs mitotic spindle organisation. Such changes suggest a role for TbATR and R-loops at trypanosome centromeres to guide kinetochore activity and chromosomes segregation (Fig.9). Loss of this activity would readily explain the profound changes in chromosome segregation we describe after TbATR RNAi. In human cells, ATR is recruited to centromeres through association with Aurora A kinase and the kinetochore component CENP-F^26^. Once recruited, ATR is activated by RPA-coated R-loops, which are generated by centromeric RNA Pol II transcription and lead to activation of Aurora B to ensure faithful chromosome segregation. A similar pathway may operate in *T. brucei*: our data suggest TbATR RNAi leads to accumulation of DNA lesions and reduced R-loop levels at the centromeres, perhaps suggesting that loss of TbATR monitoring of R-loops impedes the normal route of Aurora kinase regulation of kinetochore attachment, resulting in centromere damage (Fig.9). In support of this model, we previously described a role for TbAUK2 (Aurora A homologue) in DNA repair and in maintaining the nuclear spindle in BSF *T. brucei*^45^. However, there are several complications in comparing the role of ATR and R-loops in centromere and kinetochore function between *T. brucei* and mammals. First, though R-loops form on *T. brucei* centromeres, it is unclear if they arise from transcription at these loci. Second, the kinetoplastid kinetochore is considerably structurally diverged^92^, with no evidence that CENP-F is present, meaning it is possible that TbATR recruitment to the kinetochore acts only via TbAUK2 or through TbAUK2 association with a distinct factor. Finally, given the diverged kinetochore, it is unclear if conserved signalling pathway(s) co-ordinate chromosome segregation: three apparently kinetoplastid-specific kinase components of the kinetochore have roles in assembly and activity of the complex^93,94^; and TbAUK1 (Aurora kinase B homologue) has been implicated in directing TbATR-independent metaphase arrest via the outer kinetochore factor KKIP5 in the presence of DNA damage^95^. More recently, a further role for R-loop-directed ATR centromere activity in human cells has been described, where loss of the histone H3 variant CENP-A in S phase led to accumulation of centromeric R loops due to increased transcription, which interfered with DNA replication and lead to ATR recruitment and generated centromere damage and instability^96^. Thus, it is possible that the yH2A accumulation we detect at *T. brucei* centromeres after TbATR RNAi is connected to DNA replication and is unrelated to impairment in spindle attachment (Fig.9). Such a scenario may be consistent with early replication of the *T. brucei* centromeres^88^ but is complicated by lack of evidence that kinetoplastids encode a CENP-A homologue.

**Figure 9.**
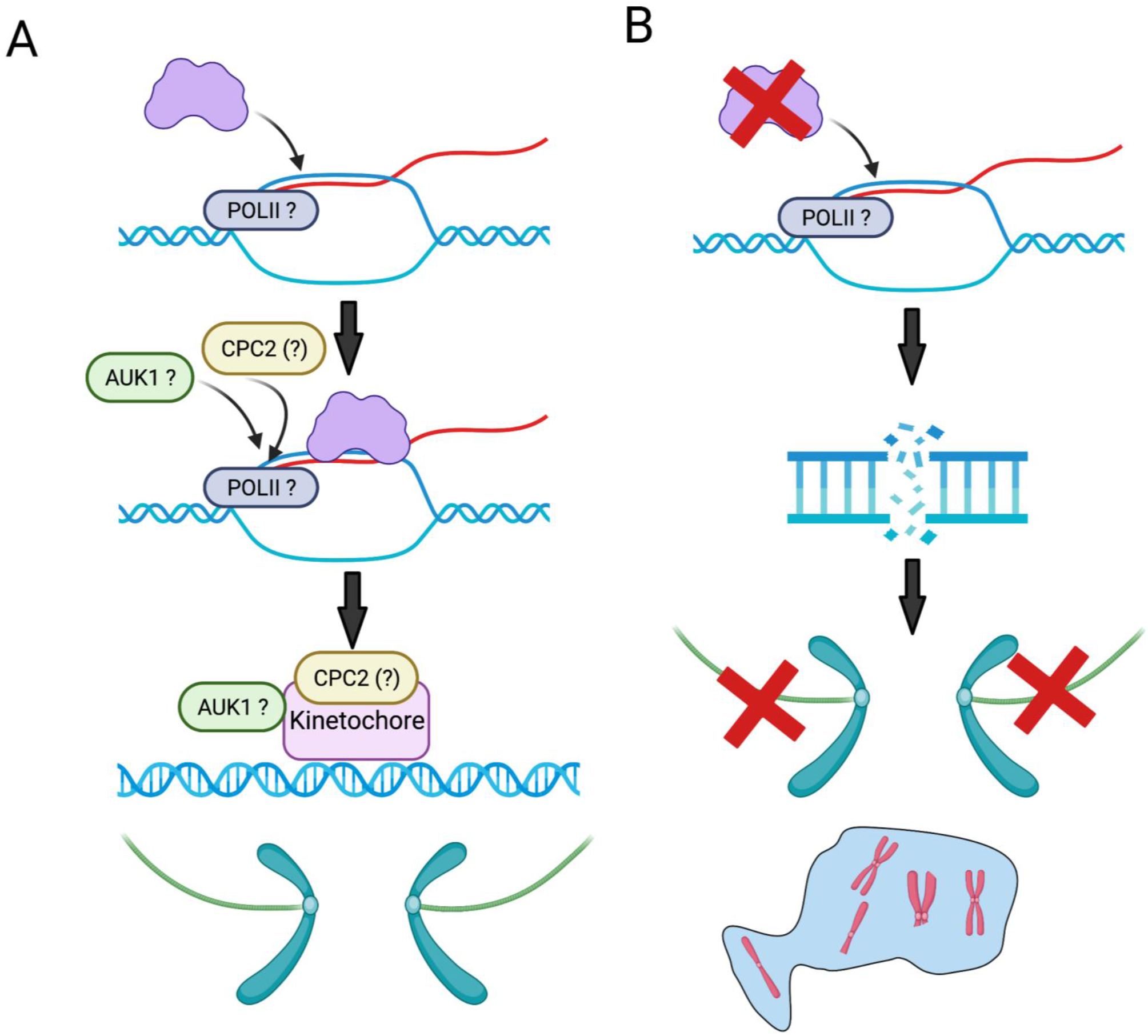
ATR acts at R-loops in *T. brucei* centromeres during chromosome segregation. (A) In the presence of TbATR, R-loop formation can be detected and resolved. What causes the formation of R-loops in the centromere is unknown but may be due to RNA Polymerase II activity. Recruitment of TbATR to centromeric R-loops may initiate activities of AUK1 (and AUK2, not shown) to ensure correct chromosome segregation via spindle attachment to the kinetochore. (B) In the absence of TbATR, R-loops in the centromere lead to DNA lesions, such as a DNA double strand break, impeding kinetochore assembly and causing chromosomes to fragment and/or segregate improperly. In turn, incorrectly segregated or fragmented chromosomes (in red) may yield aberrant nuclei, such as with a nuclear ‘bleb’.

In summary, we show widespread functions of TbATR in genome preservation and transmission in *T. brucei*, including evidence that the kinase acts upon R-loops during both gene expression and chromosome segregation, most likely explaining why TbATR is essential. Further roles of TbATR may yet emerge. In this regard it is perhaps worth noting that our analysis of chromosome distribution after RNAi relied on Telo-FISH, meaning the segregation defects we detect may extend to intermediate and minichromosomes^97^, which are a class of chromosomes peculiar to trypanosomes and in which centromeres have not been detected^98^, with mode of segregation during mitosis that appears distinct from the megabase chromosomes^99^.

## Acknowledgments

This work was supported by the BBSRC [BB/K006495/1, BB/M028909/1, BB/N016165/1 and a DTP studentship to JAB], and the MRC [MR/S019472/1]. The Wellcome Centre for Integrative Parasitology is supported by core funding from the Wellcome Trust [104111]. Glasgow Polyomics is supported by the Wellcome Trust [105614/Z/14/Z]. The Glasgow Imaging Facility is core funded by the Institute of Infection, Immunity and Inflammation (University of Glasgow). The funders had no role in study design, data collection and analysis, decision to publish, or preparation of the manuscript. We thank all members of McCulloch, Mottram and Tosi labs for fruitful discussions, support, and input into the work.

## Author contributions

JAB, JCM and RM conceived the study; JAB, KC, LL, CL, EB and RM performed the research and/or analysed the data; JAB, KC, EB, JCM, LROT and RM wrote and edited the manuscript. All authors report no conflict of interest.

## References

1. Vermeulen, K., Van Bockstaele, D. R. & Berneman, Z. N. The cell cycle: A review of regulation, deregulation and therapeutic targets in cancer. Cell Proliferation (2003). doi:10.1046/j.1365-2184.2003.00266.x

2. Sherr, C. J. & Bartek, J. Cell Cycle–Targeted Cancer Therapies. Annu. Rev. Cancer Biol. (2017). doi:10.1146/annurev-cancerbio-040716-075628

3. Minchom, A., Aversa, C. & Lopez, J. Dancing with the DNA damage response: next-generation anti-cancer therapeutic strategies. Therapeutic Advances in Medical Oncology (2018). doi:10.1177/1758835918786658

4. Saldivar, J. C. et al. An intrinsic S/G2 checkpoint enforced by ATR. Science (80-.). (2018). doi:10.1126/science.aap9346

5. Ciardo, D., Goldar, A. & Marheineke, K. On the Interplay of the DNA Replication Program and the Intra-S Phase Checkpoint Pathway. Genes 10, (2019).

6. Musacchio, A. The Molecular Biology of Spindle Assembly Checkpoint Signaling Dynamics. Current Biology (2015). doi:10.1016/j.cub.2015.08.051

7. Bartek, J. & Lukas, J. DNA damage checkpoints: from initiation to recovery or adaptation. Current Opinion in Cell Biology (2007). doi:10.1016/j.ceb.2007.02.009

8. Blackford, A. N. & Jackson, S. P. ATM, ATR, and DNA-PK: The Trinity at the Heart of the DNA Damage Response. Molecular Cell (2017). doi:10.1016/j.molcel.2017.05.015

9. Cortez, D., Guntuku, S., Qin, J. & Elledge, S. J. ATR and ATRIP: Partners in checkpoint signaling. Science (80-.). (2001). doi:10.1126/science.1065521

10. Haahr, P. et al. Activation of the ATR kinase by the RPA-binding protein ETAA1. Nat. Cell Biol. (2016). doi:10.1038/ncb3422

11. Zou, L. & Elledge, S. J. Sensing DNA damage through ATRIP recognition of RPA-ssDNA complexes. Science (80-.). (2003). doi:10.1126/science.1083430

12. Mordes, D. A., Glick, G. G., Zhao, R. & Cortez, D. TopBP1 activates ATR through ATRIP and a PIKK regulatory domain. Genes Dev. (2008). doi:10.1101/gad.1666208

13. Kumagai, A., Lee, J., Yoo, H. Y. & Dunphy, W. G. TopBP1 activates the ATR-ATRIP complex. Cell (2006). doi:10.1016/j.cell.2005.12.041

14. Delacroix, S., Wagner, J. M., Kobayashi, M., Yamamoto, K. I. & Karnitz, L. M. The Rad9-Hus1-Rad1 (9-1-1) clamp activates checkpoint signaling via TopBP1. Genes Dev. (2007). doi:10.1101/gad.1547007

15. Lee, J., Kumagai, A. & Dunphy, W. G. The Rad9-Hus1-Rad1 checkpoint clamp regulates interaction of TopBP1 with ATR. J. Biol. Chem. 282, 28036–28044 (2007).

16. Saldivar, J. C., Cortez, D. & Cimprich, K. A. The essential kinase ATR: Ensuring faithful duplication of a challenging genome. Nature Reviews Molecular Cell Biology (2017). doi:10.1038/nrm.2017.67

17. Goto, H. et al. Chk1-mediated Cdc25A degradation as a critical mechanism for normal cell cycle progression. J. Cell Sci. (2019). doi:10.1242/jcs.223123

18. Hodroj, D. et al. An ATR-dependent function for the Ddx19 RNA helicase in nuclear R-loop metabolism. EMBO J. 36, 1182–1198 (2017).

19. Matos, D. A. et al. ATR Protects the Genome against R Loops through a MUS81-Triggered Feedback Loop. Mol. Cell (2020). doi:10.1016/j.molcel.2019.10.010

20. Barroso, S. et al. The DNA damage response acts as a safeguard against harmful DNA-RNA hybrids of different origins. EMBO Rep. 20, e47250 (2019).

21. Stiff, T., Cerosaletti, K., Concannon, P., O’driscoll, M. & Jeggo, P. A. Replication independent ATR signalling leads to G2/M arrest requiring Nbs1, 53BP1 and MDC1. Hum. Mol. Genet. (2008). doi:10.1093/hmg/ddn220

22. Lin, J. J. & Dutta, A. ATR pathway is the primary pathway for activating G2/M checkpoint induction after re-replication. J. Biol. Chem. (2007). doi:10.1074/jbc.M705178200

23. Landsverk, H. B. et al. Regulation of ATR activity via the RNA polymerase II associated factors CDC73 and PNUTS-PP1. Nucleic Acids Res. 47, 1797–1813 (2019).

24. Kieft, R. et al. Identification of a novel base J binding protein complex involved in RNA polymerase II transcription termination in trypanosomes. PLOS Genet. 16, e1008390 (2020).

25. Jensen, B. C. et al. Chromatin-Associated Protein Complexes Link DNA Base J and Transcription Termination in *Leishmania*; mSphere 6, e01204–20 (2021).

26. Kabeche, L., Nguyen, H. D., Buisson, R. & Zou, L. A mitosis-specific and R loop-driven ATR pathway promotes faithful chromosome segregation. Science (80-.). (2018). doi:10.1126/science.aan6490

27. Kidiyoor, G. R., Kumar, A. & Foiani, M. ATR-mediated regulation of nuclear and cellular plasticity. DNA Repair (2016). doi:10.1016/j.dnarep.2016.05.020

28. Ma, H. & Pederson, T. The nucleolus stress response is coupled to an ATR-Chk1-mediated G2 arrest. Mol. Biol. Cell 24, 1334–1342 (2013).

29. Kidiyoor, G. R. et al. ATR is essential for preservation of cell mechanics and nuclear integrity during interstitial migration. Nat. Commun. 11, 4828 (2020).

30. Kumar, A. et al. ATR Mediates a Checkpoint at the Nuclear Envelope in Response to Mechanical Stress. Cell 158, 633–646 (2014).

31. Brown, N. & Costanzo, V. An ATM and ATR dependent pathway targeting centrosome dependent spindle assembly. Cell Cycle 8, 1997–2001 (2009).

32. Büscher, P., Cecchi, G., Jamonneau, V. & Priotto, G. Human African trypanosomiasis. The Lancet (2017). doi:10.1016/S0140-6736(17)31510-6

33. Parsons, M., Worthey, E. A., Ward, P. N. & Mottram, J. C. Comparative analysis of the kinomes of three pathogenic trypanosomatids: Leishmania major, Trypanosoma brucei and Trypanosoma cruzi. BMC Genomics (2005). doi:10.1186/1471-2164-6-127

34. Black, J. A. et al. Trypanosoma brucei ATR Links DNA Damage Signaling during Antigenic Variation with Regulation of RNA Polymerase I-Transcribed Surface Antigens. Cell Rep. (2020). doi:10.1016/j.celrep.2019.12.049

35. Jones, N. G. et al. Regulators of Trypanosoma brucei Cell Cycle Progression and Differentiation Identified Using a Kinome-Wide RNAi Screen. PLoS Pathog. 10, (2014).

36. Stortz, J. A. et al. Genome-wide and protein kinase-focused RNAi screens reveal conserved and novel damage response pathways in Trypanosoma brucei. PLOS Pathog. 13, e1006477 (2017).

37. Marin, P. A. et al. ATR Kinase Is a Crucial Player Mediating the DNA Damage Response in Trypanosoma brucei. Frontiers in Cell and Developmental Biology 8, 1651 (2020).

38. Genois, M.-M. et al. DNA Repair Pathways in Trypanosomatids: from DNA Repair to Drug Resistance. Microbiol. Mol. Biol. Rev. (2014). doi:10.1128/mmbr.00045-13

39. Damasceno, J. D. et al. Conditional genome engineering reveals canonical and divergent roles for the Hus1 component of the 9–1–1 complex in the maintenance of the plastic genome of Leishmania. Nucleic Acids Res. (2018). doi:10.1093/nar/gky1017

40. Damasceno, J. D. et al. Functional compartmentalization of Rad9 and Hus1 reveals diverse assembly of the 9-1-1 complex components during the DNA damage response in Leishmania. Mol. Microbiol. (2016). doi:10.1111/mmi.13441

41. Altschul, S. F. et al. Gapped BLAST and PSI-BLAST: A new generation of protein database search programs. Nucleic Acids Research 25, 3389–3402 (1997).

42. Kelly, S. et al. Functional genomics in Trypanosoma brucei: A collection of vectors for the expression of tagged proteins from endogenous and ectopic gene loci. Mol. Biochem. Parasitol. 154, 103–109 (2007).

43. Devlin, R. et al. Mapping replication dynamics in Trypanosoma brucei reveals a link with telomere transcription and antigenic variation. Elife 5, e12765 (2016).

44. Müller, L. S. M. et al. Genome organization and DNA accessibility control antigenic variation in trypanosomes. Nature (2018). doi:10.1038/s41586-018-0619-8

45. Stortz, J. A. et al. Genome-wide and protein kinase-focused RNAi screens reveal conserved and novel damage response pathways in Trypanosoma brucei. PLoS Pathog. (2017). doi:10.1371/journal.ppat.1006477

46. Alsford, S., Kawahara, T., Glover, L. & Horn, D. Tagging a T. brucei RRNA locus improves stable transfection efficiency and circumvents inducible expression position effects. Mol. Biochem. Parasitol. (2005). doi:10.1016/j.molbiopara.2005.08.009

47. Hirumi, H. & Hirumi, K. Continuous Cultivation of Trypanosoma brucei Blood Stream Forms in a Medium Containing a Low Concentration of Serum Protein without Feeder Cell Layers. J. Parasitol. 75, 985–989 (1989).

48. Glover, L., Marques, C. A., Suska, O. & Horn, D. Persistent DNA damage Foci and DNA replication with a broken chromosome in the African trypanosome. MBio (2019). doi:10.1128/mBio.01252-19

49. Schindelin, J. et al. Fiji: an open-source platform for biological-image analysis. Nat Meth 9, 676–682 (2012).

50. Schindelin, J. et al. Fiji: An open-source platform for biological-image analysis. Nature Methods (2012). doi:10.1038/nmeth.2019

51. Alexa, A., Rahnenführer, J. & Lengauer, T. Improved scoring of functional groups from gene expression data by decorrelating GO graph structure. Bioinformatics 22, 1600–1607 (2006).

52. Briggs, E., Hamilton, G., Crouch, K., Lapsley, C. & McCulloch, R. Genome-wide mapping reveals conserved and diverged R-loop activities in the unusual genetic landscape of the African trypanosome genome. Nucleic Acids Res. (2018). doi:10.1093/nar/gky928

53. Langmead, B. & Salzberg, S. L. Fast gapped-read alignment with Bowtie 2. Nat. Methods (2012). doi:10.1038/nmeth.1923

54. Li, H. et al. The Sequence Alignment/Map format and SAMtools. Bioinformatics (2009). doi:10.1093/bioinformatics/btp352

55. Ramírez, F., Dündar, F., Diehl, S., Grüning, B. A. & Manke, T. DeepTools: A flexible platform for exploring deep-sequencing data. Nucleic Acids Res. (2014). doi:10.1093/nar/gku365

56. Diaz, A., Park, K., Lim, D. A. & Song, J. S. Normalization, bias correction, and peak calling for ChIP-seq. Stat. Appl. Genet. Mol. Biol. (2012). doi:10.1515/1544-6115.1750

57. Robinson, J. T. et al. Integrative genomics viewer. Nature Biotechnology (2011). doi:10.1038/nbt.1754

58. Hahne, F. & Ivanek, R. Statistical genomics: methods and protocols. Chapter Vis. genomic data using Gviz Bioconductor. New York Springer New York (2016).

59. Afgan, E. et al. The Galaxy platform for accessible, reproducible and collaborative biomedical analyses: 2018 update. Nucleic Acids Res. (2018). doi:10.1093/nar/gky379

60. Wickham, H. ggplot2 Elegant Graphics for Data Analysis. Journal of the Royal Statistical Society: Series A (Statistics in Society) (2016). doi:10.1007/978-3-319-24277-4

61. DuBois, K. N. et al. NUP-1 is a large coiled-coil nucleoskeletal protein in trypanosomes with lamin-like functions. PLoS Biol. (2012). doi:10.1371/journal.pbio.1001287

62. Glover, L. & Horn, D. Trypanosomal histone γh2A and the DNA damage response. Mol. Biochem. Parasitol. (2012). doi:10.1016/j.molbiopara.2012.01.008

63. Ivens, A. C. et al. The genome of the kinetoplastid parasite, Leishmania major. Science (80-.). (2005). doi:10.1126/science.1112680

64. Cestari, I. & Stuart, K. Inositol phosphate pathway controls transcription of telomeric expression sites in trypanosomes. Proc. Natl. Acad. Sci. 112, E2803 LP–E2812 (2015).

65. Leal, A. Z. et al. Genome maintenance functions of a putative Trypanosoma brucei translesion DNA polymerase include telomere association and a role in antigenic variation. Nucleic Acids Res. 48, 9660–9680 (2020).

66. Briggs, E. et al. Trypanosoma brucei ribonuclease H2A is an essential R-loop processing enzyme whose loss causes DNA damage during transcription initiation and antigenic variation. Nucleic Acids Res. (2019). doi:10.1093/nar/gkz644

67. Briggs, E., Crouch, K., Lemgruber, L., Lapsley, C. & McCulloch, R. Ribonuclease H1-targeted R-loops in surface antigen gene expression sites can direct trypanosome immune evasion. PLoS Genet. (2018). doi:10.1371/journal.pgen.1007729

68. Tu, X., Kumar, P., Li, Z. & Wang, C. C. An Aurora Kinase Homologue Is Involved in Regulating Both Mitosis and Cytokinesis in <em>Trypanosoma brucei</em>*. J. Biol. Chem. 281, 9677–9687 (2006).

69. Lamm, N. et al. Nuclear F-actin counteracts nuclear deformation and promotes fork repair during replication stress. Nat. Cell Biol. 22, 1460–1470 (2020).

70. Li, Z. & Wang, C. C. Changing roles of aurora-B kinase in two life cycle stages of Trypanosoma brucei. Eukaryot. Cell 5, 1026–1035 (2006).

71. Da Silva, R. B., Machado, C. R., Aquiles Rodrigues, A. R. & Pedrosa, A. L. Selective human inhibitors of ATR and ATM render leishmania major promastigotes sensitive to oxidative damage. PLoS One (2018). doi:10.1371/journal.pone.0205033

72. Sancar, A., Lindsey-Boltz, L. A., Unsal-Kaçmaz, K. & Linn, S. Molecular mechanisms of mammalian DNA repair and the DNA damage checkpoints. Annu. Rev. Biochem. 73, 39–85 (2004).

73. Damasceno, J. D. et al. Genome duplication in leishmania major relies on persistent subtelomeric dna replication. Elife (2020). doi:10.7554/ELIFE.58030

74. Moiseeva, T. N. et al. An ATR and CHK1 kinase signaling mechanism that limits origin firing during unperturbed DNA replication. Proc. Natl. Acad. Sci. 116, 13374 LP–13383 (2019).

75. Rainey, M. D. et al. ATR Restrains DNA Synthesis and Mitotic Catastrophe in Response to CDC7 Inhibition. Cell Rep. 32, 108096 (2020).

76. Ibarra, A., Schwob, E. & Méndez, J. Excess MCM proteins protect human cells from replicative stress by licensing backup origins of replication. Proc. Natl. Acad. Sci. 105, 8956 LP–8961 (2008).

77. Woodward, A. M. et al. Excess Mcm2-7 license dormant origins of replication that can be used under conditions of replicative stress. J. Cell Biol. 173, 673–683 (2006).

78. Calderano, S. G. et al. Single molecule analysis of Trypanosoma brucei DNA replication dynamics. Nucleic Acids Res. 43, 2655–2665 (2015).

79. Lee, K. J. & Li, Z. The CRK2-CYC13 complex functions as an S-phase cyclin-dependent kinase to promote DNA replication in Trypanosoma brucei. BMC Biol. 19, 29 (2021).

80. Damasceno, J. D., Marques, C. A., Black, J., Briggs, E. & McCulloch, R. Read, Write, Adapt: Challenges and Opportunities during Kinetoplastid Genome Replication. Trends Genet. 37, 21–34 (2021).

81. Hamperl, S., Bocek, M. J., Saldivar, J. C., Swigut, T. & Cimprich, K. A. Transcription-Replication Conflict Orientation Modulates R-Loop Levels and Activates Distinct DNA Damage Responses. Cell 170, 774-786.e19 (2017).

82. Shao, X., Joergensen, A. M., Howlett, N. G., Lisby, M. & Oestergaard, V. H. A distinct role for recombination repair factors in an early cellular response to transcription-replication conflicts. Nucleic Acids Res. 48, 5467–5484 (2020).

83. Helmrich, A., Ballarino, M. & Tora, L. Collisions between replication and transcription complexes cause common fragile site instability at the longest human genes. Mol. Cell 44, 966–977 (2011).

84. Pentzold, C. et al. FANCD2 binding identifies conserved fragile sites at large transcribed genes in avian cells. Nucleic Acids Res. 46, 1280–1294 (2018).

85. Wilson, T. E. et al. Large transcription units unify copy number variants and common fragile sites arising under replication stress. Genome Res. 25, 189–200 (2015).

86. Daniels, J.-P., Gull, K. & Wickstead, B. Cell biology of the trypanosome genome. Microbiol. Mol. Biol. Rev. 74, 552–569 (2010).

87. da Silva, M. S. et al. Transcription activity contributes to the firing of non-constitutive origins in African trypanosomes helping to maintain robustness in S-phase duration. Sci. Rep. 9, 18512 (2019).

88. Tiengwe, C. et al. Genome-wide analysis reveals extensive functional interaction between DNA replication initiation and transcription in the genome of trypanosoma brucei. Cell Rep. (2012). doi:10.1016/j.celrep.2012.06.007

89. Teloni, F. et al. Efficient Pre-mRNA Cleavage Prevents Replication-Stress-Associated Genome Instability. Mol. Cell 73, 670-683.e12 (2019).

90. Matthews, K. R., Tschudi, C. & Ullu, E. A common pyrimidine-rich motif governs trans-splicing and polyadenylation of tubulin polycistronic pre-mRNA in trypanosomes. Genes Dev. 8, 491–501 (1994).

91. Devlin, R., Marques, C. A. & McCulloch, R. Does DNA replication direct locus-specific recombination during host immune evasion by antigenic variation in the African trypanosome? Curr. Genet. 63, 441–449 (2017).

92. Akiyoshi, B. & Gull, K. Discovery of unconventional kinetochores in kinetoplastids. Cell (2014). doi:10.1016/j.cell.2014.01.049

93. Ishii, M. & Akiyoshi, B. Characterization of unconventional kinetochore kinases KKT10/19 in Trypanosoma brucei. J. Cell Sci. jcs.240978 (2020). doi:10.1242/jcs.240978

94. Saldivia, M. et al. Targeting the trypanosome kinetochore with CLK1 protein kinase inhibitors. Nat. Microbiol. 5, 1207–1216 (2020).

95. Zhou, Q., Pham, K. T. M., Hu, H., Kurasawa, Y. & Li, Z. A kinetochore-based ATM/ATR-independent DNA damage checkpoint maintains genomic integrity in trypanosomes. Nucleic Acids Res. (2019). doi:10.1093/nar/gkz476

96. Giunta, S. et al. CENP-A chromatin prevents replication stress at centromeres to avoid structural aneuploidy. Proc. Natl. Acad. Sci. 118, e2015634118 (2021).

97. Wickstead, B., Ersfeld, K. & Gull, K. The mitotic stability of the minichromosomes of Trypanosoma brucei. Mol. Biochem. Parasitol. (2003). doi:10.1016/j.molbiopara.2003.08.007

98. Wickstead, B., Ersfeld, K. & Gull, K. The small chromosomes of Trypanosoma brucei involved in antigenic variation are constructed around repetitive palindromes. Genome Res. 14, 1014–1024 (2004).

99. Ersfeld, K. & Gull, K. Partitioning of Large and Minichromosomes in *Trypanosoma brucei*; Science (80-.). 276, 611 LP–614 (1997).

