## Supplementary Figures for "Widespread roles of *Trypanosoma brucei* ATR in nuclear genome function and transmission are linked to R-loops"

### Supporting Information and Supplementary Figures

**Table S1.** Underlying data for figures.

**Table S2.** Reagents and oligonucleotides used in this study.

**Table S3.** RNA-seq data for all genes significantly up regulated at 36 hrs post induction.

**Table S4.** RNA-seq data for all genes significantly downregulated at 36 hrs post induction.

**Supplementary File 1. Whole chromosome plots of yH2A ChIP-seq and DRIP-seq signal.** The centromere position is marked by a red line and CDSs are marked as red (-ve strand) or grey (+ve strand) arrows. The rRNA locus is marked in green.

### Supplementary Figure Legends

**Figure S1. Electron microscopy of cells after TbATR RNAi.** (A) Representative images of TbATR RNAi CL1 cells after 36 hrs growth without RNAi induction. Nuclear and kinetoplast DNA is stained with DAPI (cyan) and the cell morphology visualised by DIC imaging. Scale bar = 10 μm. (B) Representative Transmission EM (TEM) images of cells in the absence (left, Tet-) or presence (Tet+) of TbATR RNAi after 24 hrs. Scale bars are as detailed on the images. (C) Representative TEM images of cells in the absence (top left, Tet-) or presence (Tet+) of TbATR RNAi induction after 36 hrs. Black arrow indicates a nuclear 'bleb', white arrows indicate additional nuclear membranes, black stars highlight 3 separate flagella associated with the flagellar pocket (the site of endo- and exo-cytosis in T. *brucei*).

**Figure S2. Cells continue to replicate in after TbATR RNAi.** (A) Representative images of 2T1 cells (the RNAi parental cell line) after 36 hrs growth with (T+) and without tetracycline (T-) and stained for EdU incorporation and probed for yH2A signal. Nuclear and kinetoplast DNA were stained with DAPI (cyan), EdU shown in red and yH2A signal green. The cell body was visualised by DIC imaging. Scale bars = 10 μm. (B) Representative field of view (FOV) images of TbATR RNAi CL1 cells at 36 hrs +/- tetracycline after EdU and yH2A labelling (shown as above). Scale bars = 20 μm.

**Figure S3. TbATR RNAi induced cells are defective in chromosome segregation.** (A). Representative super resolution images of telomere distribution in the nuclei of uninduced TbATR RNAi cells after 36 hrs growth. Images were captured and visualised using SR-SIM microscopy on an Elyra SR microscope (Zeiss). Scale bar = 2 μm. The nuclear and kinetoplast DNA were visualised with DAPI (magenta) and the telomeres by a telomeric FITC conjugated probe (green). (B) Representative super resolution images of telomeric distribution in the nuclei of TbATR depleted RNAi cells after 36 hrs induction. Images were captured and are shown as in (A). (C) Representative field of view images after 36 hrs growth in the presence (+) or absence (-) of TbATR RNAi. Scale bars = 10 μm.

**Figure S4. Significantly up or down regulated GO terms following depletion of TbATR by RNAi at 36 hrs** (A) Up-regulated and (B) down-regulated GO terms were categorised as Biological Process (BP) or Metabolic Function (MF). Pie charts show the distribution of significantly changed GO terms (as a percentage of the total number of BP or MF associated GO terms) following a Fisher's Exact test. $p < 0.05$.

**Figure S5. R-loop and yH2A distribution across transcription initiation and termination sites in the genome of *T. brucei* BSF cells.** Metaplots and heatmaps show (A) yH2A signal mapped across strand switch regions (SSRs). SSR regions are scaled to 500 bp and +/- 2.0 Kb is plotted around them. dTSS (divergent transcription start site), cTTS (convergent transcription termination site), head-to-tail regions (sTTS [split transcription termination site], sTSS [split transcription start site]). (B) Metaplots and heatmaps show and DRIPseq (R-loop) signal mapped across strand switch regions (SSRs). SSR regions are scaled to 1 kb bp and +/- 10 Kb is plotted around them. Labels are as stated in (A).

**Figure S6. yH2A ChIPseq and DRIPseq signal mapped across all silent BESs after TbATR RNAi.** Whole BES plots show yH2A signal enrichment (normalised to the corresponding input and subsequently normalised to the uninduced sample) and DRIPseq signal (normalised to the corresponding input controls). Signal was mapped using Gviz. BESs are numbered and genes are as indicated in Fig.5.

**Figure S7. yH2A ChIPseq and DRIPseq signal mapped across all centromeres after TbATR RNAi.** yH2A and DRIPseq signal plotted across all centromeric regions; data are plotted as shown in Fig.S6. The centromere position is marked by a black box and a centromeric proximal CDS (centromere 3, in chromosome 3) is marked by a grey arrow (+ve strand).

**Figure S8. TbATR depletion is correlates with deficient mitotic spindle formation 24 hrs after RNAi induction.** (A) Representative images of antibody negative control cells. The nuclear and kinetoplast DNA were stained with DAPI (grey). Images were captured on an Axioskop 2 (Zeiss). Scale bars = 10 μm. (B) Cells were collected at 24 hrs post induction of TbATR, and the mitotic spindle detected via indirect immunofluorescence (as described in Fig.7) and the number cells in the population containing a recognisable mitotic spindle were counted and represented as a perecentage of toral cells. Error bars = ± SEM, n=3 independent experiments (>100 cells counted/experiment). (*) p = 0.0316, unpaired t-test (two-tailed).

**Figure S9. Validation of cells expressing TbATR endogenously 12 myc tagged at the N-terminus ($^{12myc}$ATR+/-).** (A) Schematic cartoon of the strategy used to endogenously tag TbATR at the N-terminus and to delete one allele generating a heterozygote, tagged cell line. Positions of primers used for cell line confirmation are marked by black arrows. (B) PCR analysis of $^{12myc}$ATR+/- clones. Clone C4 was selected for further analysis. DNA ladder sizes are in bp. (C) Western blot analysis confirming expression of endogenously myc-tagged TbATR. The predicted size of myc-tagged TbATR is 320.594 kD (ATR) + 14.4 kDa (12xmyc epitope; total = ~334.994 kDa. (*) marks the selected clone (C4). (D) Growth curve analysis of WT, $^{12myc}$ATR+/+ and $^{12myc}$ATR+/- cell lines in the presence and absence of 0.0003% MMS. MMS stock was prepared in HMI-11, 20% FBS. Cell density was assessed every 24 hrs for 72 hrs. Error bars = ± SEM, n=3 independent experiments. (E) The intensities of DAPI (nDNA; grey) and myc signal ($^{12myc}$ATR; red) were measured using Fiji (ImageJ). A region of interest (21x21 pixels) was drawn around individual nuclei. The mean pixel was then measured. Each point on the graph represents signal from a nucleus. The errors bars represent the mean value and the standard deviation. A Kruskal-Wallis non-parametric test was used to assess significance. (****) p = <0.0001, (***) p = 0.0001, ns = non-significant. Data from one experiment (n = 227 nuclei). Background intensity was calculated using WT untagged cells (324.831 A.U.; arbitrary units; dashed line). (F) The intensities of DAPI (nDNA; grey) and myc signal ($^{12myc}$ATR; red) were measured using Fiji (ImageJ). A region of interest (21x21 pixels) was drawn around individual nuclei. The mean pixel was then measured. Each point on the graph represents signal from a nucleus. The errors bars represent the mean value and the standard deviation. A Kruskal-Wallis non-parametric test was used to assess significance. DAPI (****) p = <0.0001, (*) p = 0.0499, FITC (***) p = equal to or >0.0001, (*) p = 0.0108, ns = non-significant. Data from one experiment (n = 156 nuclei). Background intensity was calculated using WT untagged cells (79.59 A.U.; arbitrary units; dashed line). (G) Additional representative images of the sub-cellular localisation of the myc signal in $^{12myc}$ATR+/- mitotic cells. White arrows mark the thread-like myc signal. Images were captured on DeltaVision Core microscope (AppliedPrecision). Nuclear and kinetoplast DNA (cyan), myc signal (yellow). Cell bodies were visualised by brightfield microscopy. Scale bar = 10 μm.

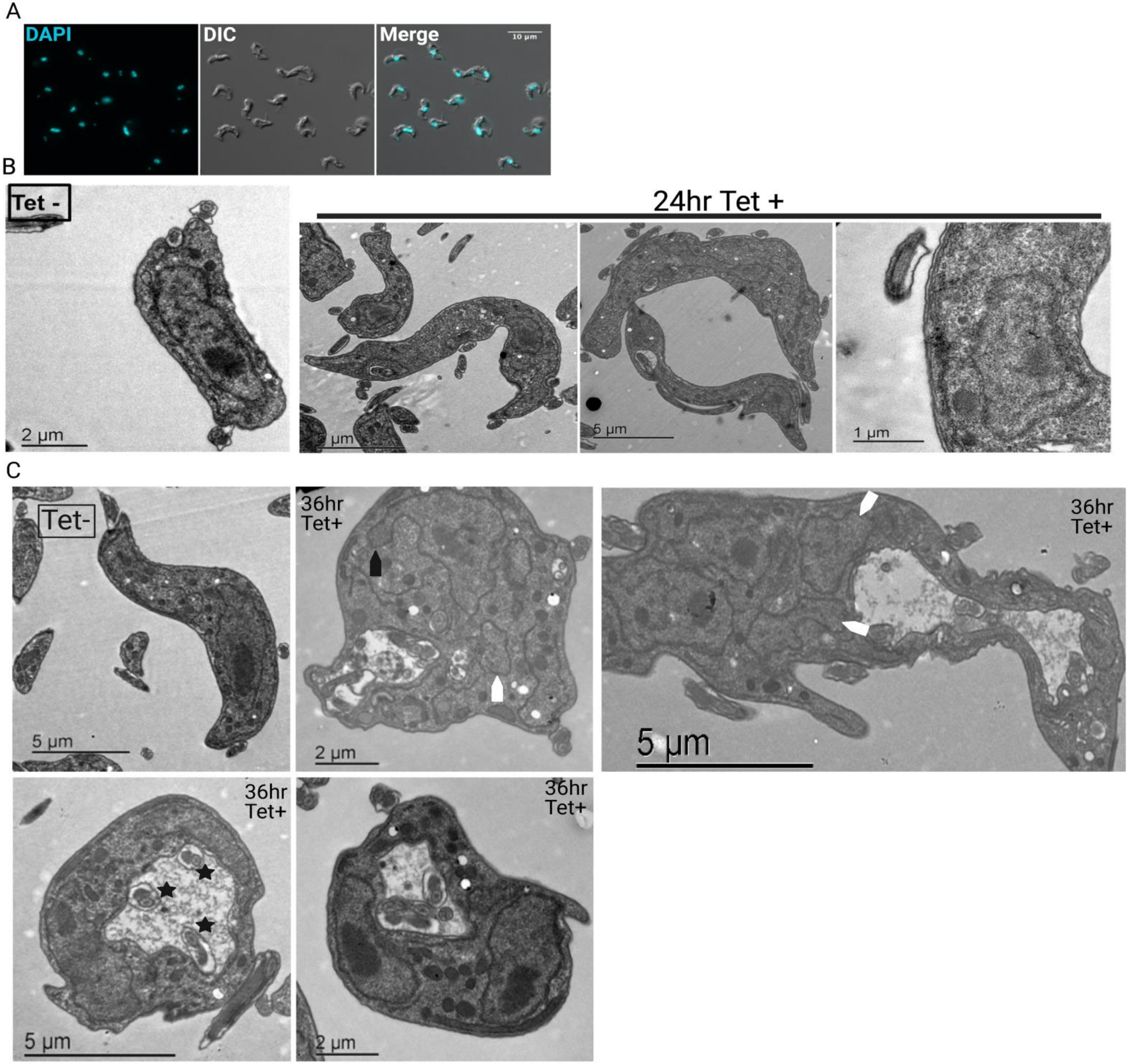

A
DAPI
DIC
Merge
10 µm
B
Tet -
24hr Tet +
2 µm
5 µm
5 µm
1 µm
C
Tet-
36hr Tet+
36hr Tet+
5 µm
2 µm
5 µm
36hr Tet+
36hr Tet+
5 µm
2 µm


**Figure S1**

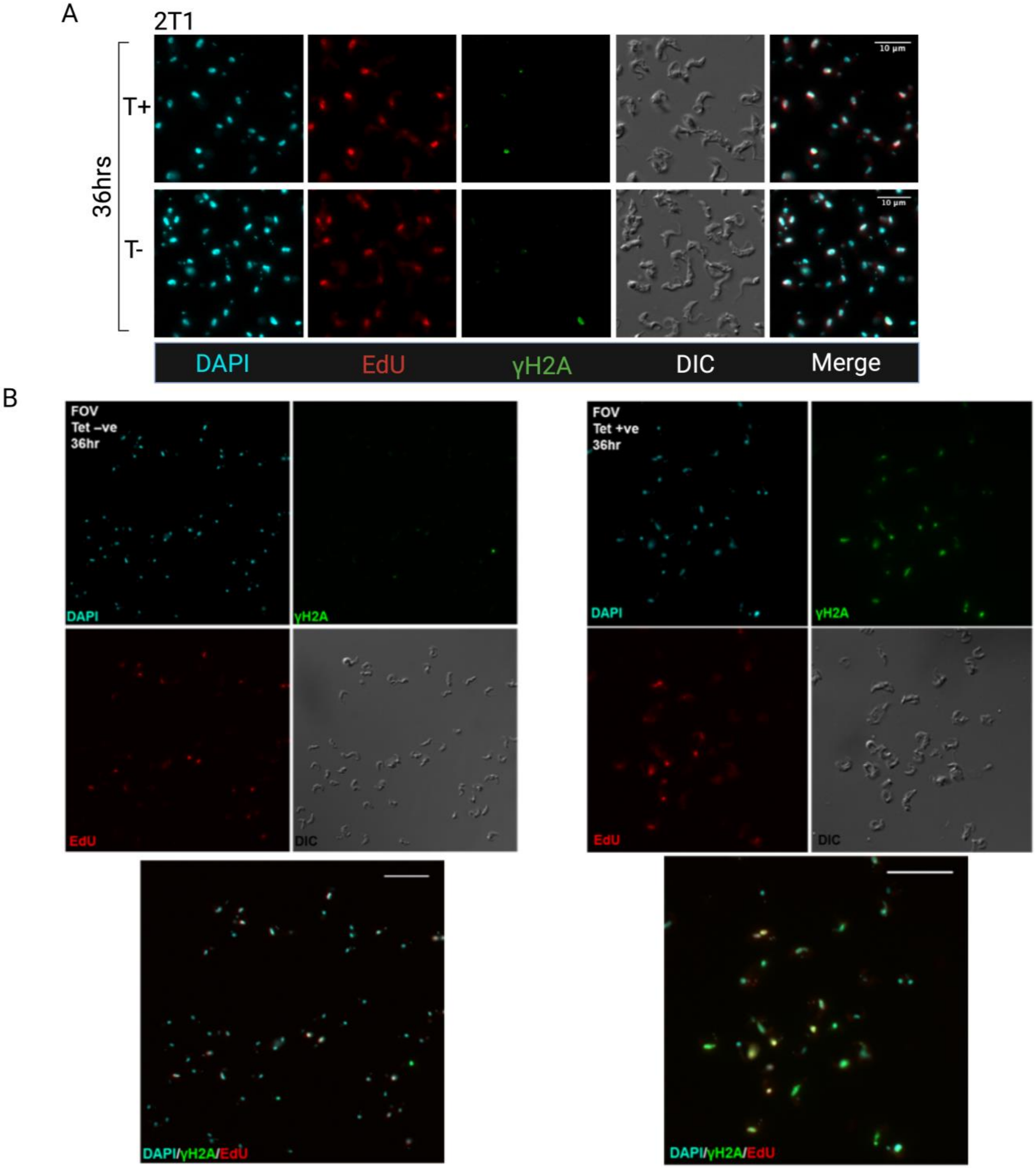
A
2T1
36hrs
T+
T-
10 µm
DAPI
EdU
γH2A
DIC
Merge
B
FOV
Tet –ve
36hr
DAPI
γH2A
EdU
DIC
DAPI/γH2A/EdU
FOV
Tet +ve
36hr
DAPI
γH2A
EdU
DIC
DAPI/γH2A/EdU

Figure S2

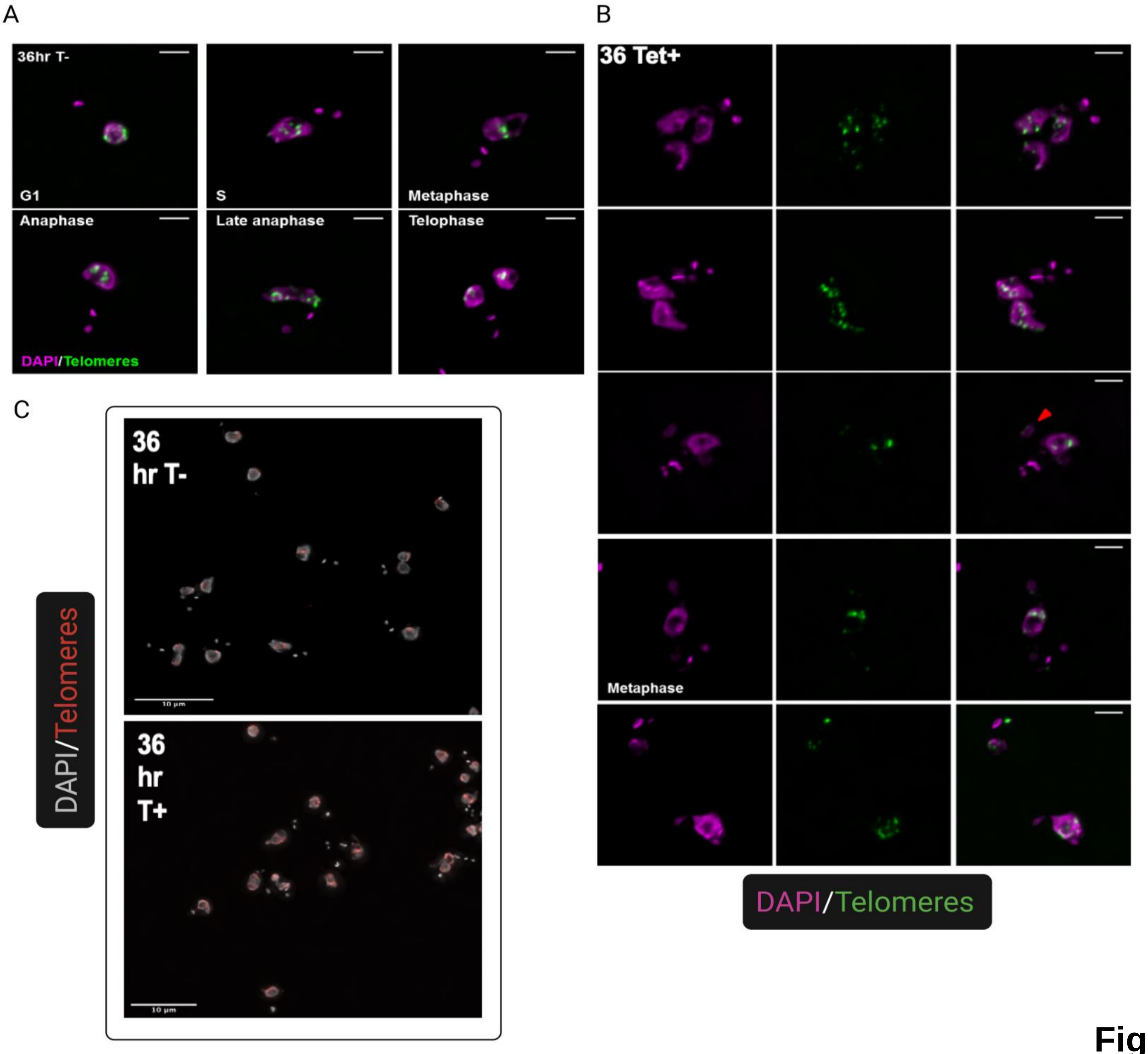
A
36hr T-
G1
S
Metaphase
Anaphase
Late anaphase
Telophase
DAPI/Telomeres
B
36 Tet+
Metaphase
DAPI/Telomeres
C
36 hr T-
36 hr T+
DAPI/Telomeres

**Figure S3**

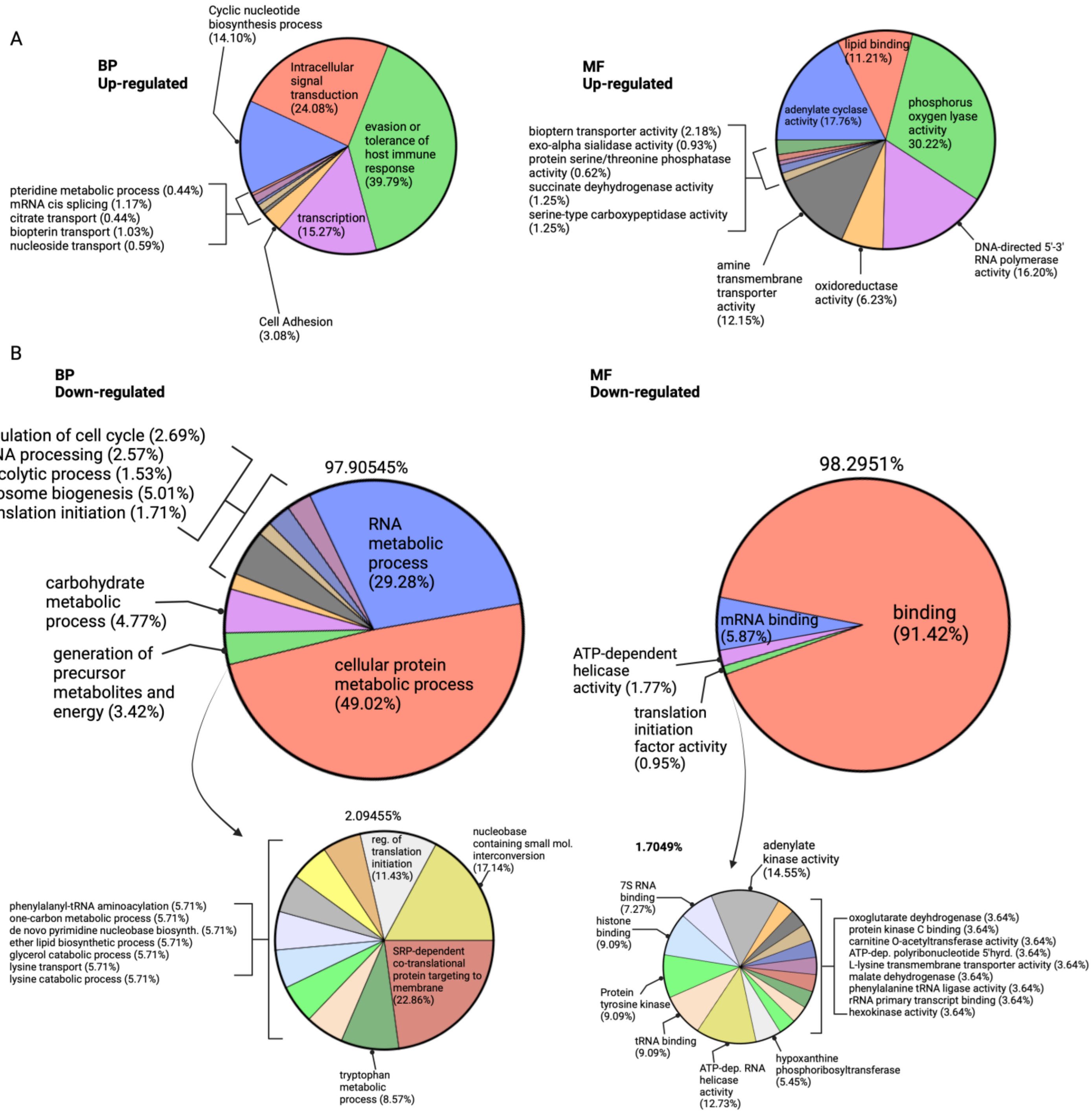
A
BP
Up-regulated
Cyclic nucleotide biosynthesis process (14.10%)
Intracellular signal transduction (24.08%)
evasion or tolerance of host immune response (39.79%)
transcription (15.27%)
pteridine metabolic process (0.44%)
mRNA cis splicing (1.17%)
citrate transport (0.44%)
biopterin transport (1.03%)
nucleoside transport (0.59%)
Cell Adhesion (3.08%)
MF
Up-regulated
lipid binding (11.21%)
adenylate cyclase activity (17.76%)
phosphorus oxygen lyase activity 30.22%)
biopterin transporter activity (2.18%)
exo-alpha sialidase activity (0.93%)
protein serine/threonine phosphatase activity (0.62%)
succinate deyhydrogenase activity (1.25%)
serine-type carboxypeptidase activity (1.25%)
amine transmembrane transporter activity (12.15%)
oxidoreductase activity (6.23%)
DNA-directed 5'-3' RNA polymerase activity (16.20%)
B
BP
Down-regulated
regulation of cell cycle (2.69%)
rRNA processing (2.57%)
glycolytic process (1.53%)
ribosome biogenesis (5.01%)
translation initiation (1.71%)
97.90545%
RNA metabolic process (29.28%)
carbohydrate metabolic process (4.77%)
generation of precursor metabolites and energy (3.42%)
cellular protein metabolic process (49.02%)
MF
Down-regulated
98.2951%
binding (91.42%)
mRNA binding (5.87%)
ATP-dependent helicase activity (1.77%)
translation initiation factor activity (0.95%)
2.09455%
reg. of translation initiation (11.43%)
nucleobase containing small mol. interconversion (17.14%)
phenylalanyl-tRNA aminoacylation (5.71%)
one-carbon metabolic process (5.71%)
de novo pyrimidine nucleobase biosynth. (5.71%)
ether lipid biosynthetic process (5.71%)
glycerol catabolic process (5.71%)
lysine transport (5.71%)
lysine catabolic process (5.71%)
SRP-dependent co-translational protein targeting to membrane (22.86%)
tryptophan metabolic process (8.57%)
1.7049%
adenylate kinase activity (14.55%)
7S RNA binding (7.27%)
histone binding (9.09%)
Protein tyrosine kinase (9.09%)
tRNA binding (9.09%)
ATP-dep. RNA helicase activity (12.73%)
hypoxanthine phosphoribosyltransferase (5.45%)
oxoglutarate deyhdrogenase (3.64%)
protein kinase C binding (3.64%)
carnitine O-acetyltransferase activity (3.64%)
ATP-dep. polyribonucleotide 5'hyrd. (3.64%)
L-lysine transmembrane transporter activity (3.64%)
malate dehydrogenase (3.64%)
phenylalanine tRNA ligase activity (3.64%)
rRNA primary transcript binding (3.64%)
hexokinase activity (3.64%)

**Figure S4**

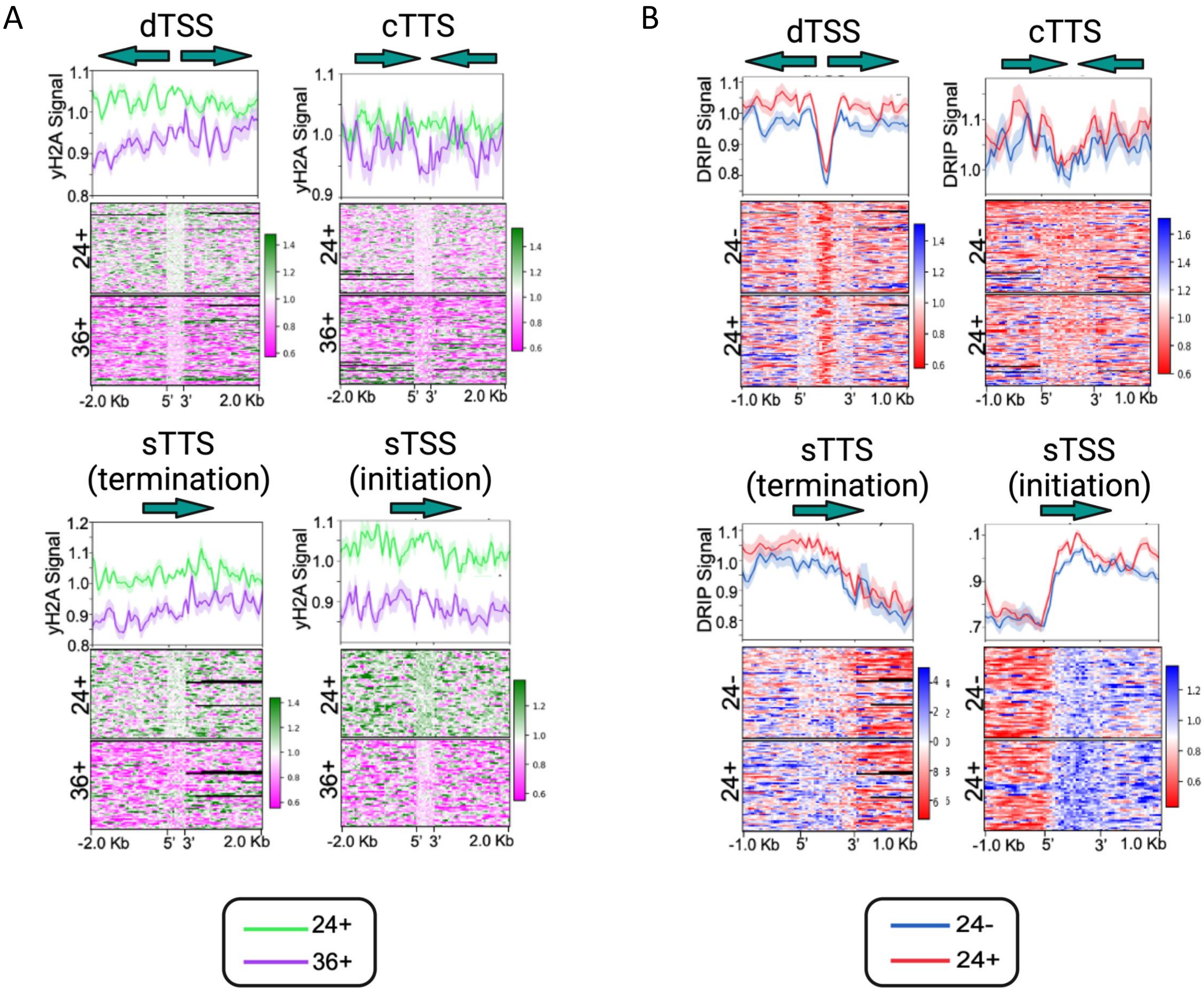
A
dTSS
cTTS
yH2A Signal
24+
36+
-2.0 Kb
5' 3'
2.0 Kb
sTTS
(termination)
sTSS
(initiation)
B
DRIP Signal
24-
-1.0 Kb
1.0 Kb
24+
36+
24-
24+

**Figure S5**

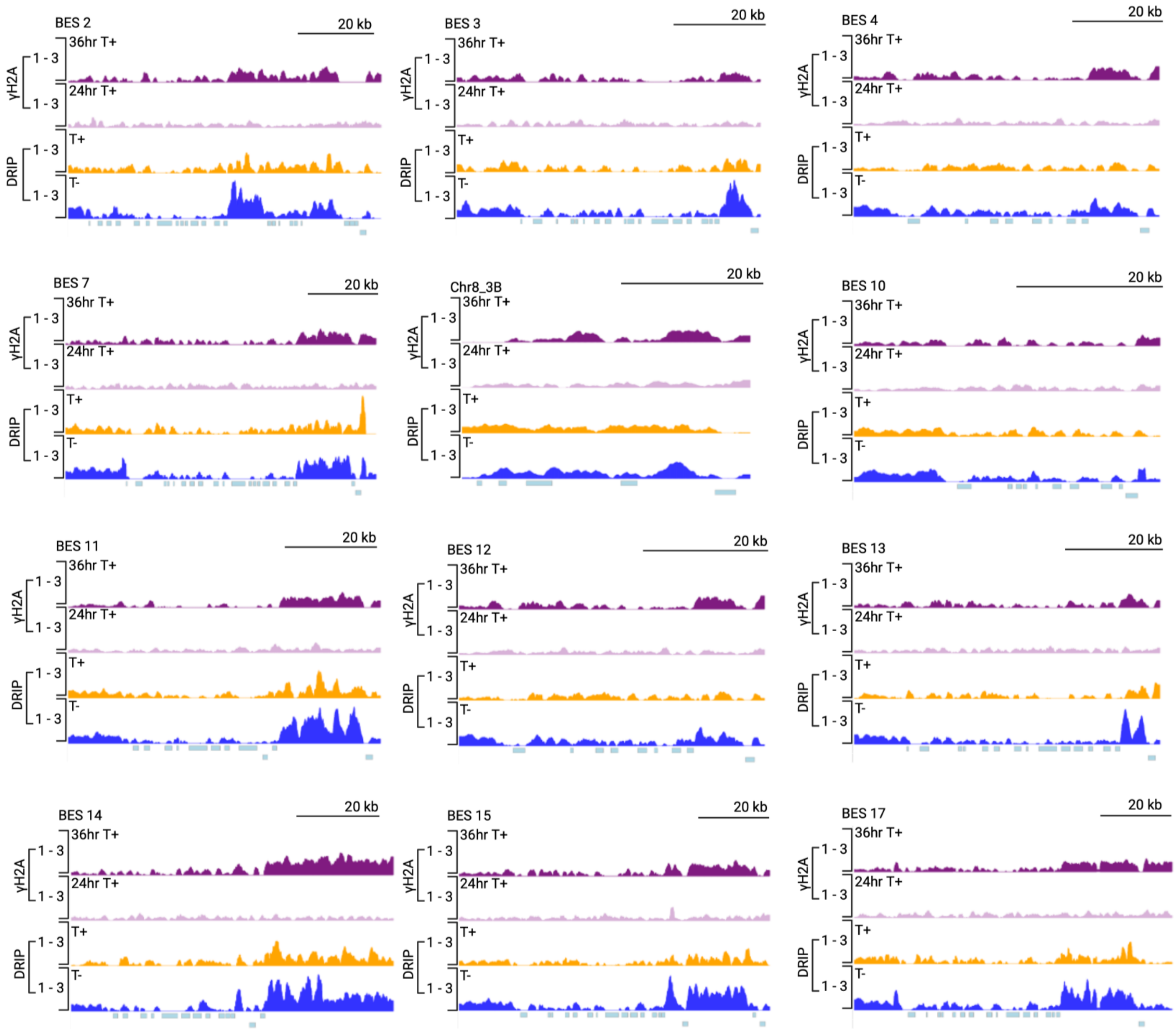
BES 2
20 kb
BES 3
20 kb
BES 4
20 kb
BES 7
20 kb
Chr8_3B
20 kb
BES 10
20 kb
BES 11
20 kb
BES 12
20 kb
BES 13
20 kb
BES 14
20 kb
BES 15
20 kb
BES 17
20 kb
γH2A
DRIP
36hr T+
24hr T+
T+
T-
1 - 3

**Figure S6**

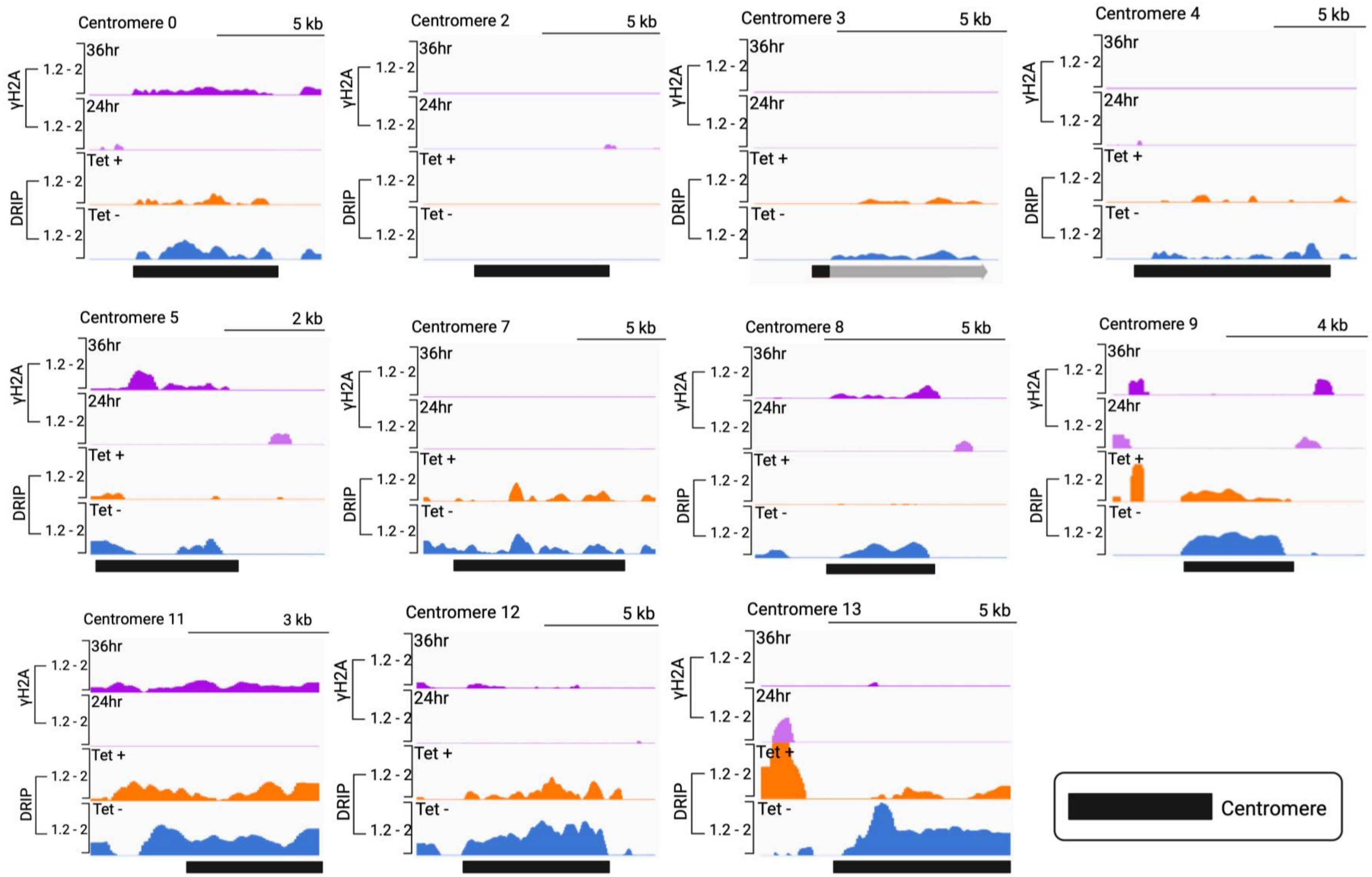
Centromere 0
5 kb
Centromere 2
5 kb
Centromere 3
5 kb
Centromere 4
5 kb
Centromere 5
2 kb
Centromere 7
5 kb
Centromere 8
5 kb
Centromere 9
4 kb
Centromere 11
3 kb
Centromere 12
5 kb
Centromere 13
5 kb
γH2A
DRIP
36hr
24hr
Tet +
Tet -
1.2 - 2
Centromere

**Figure S7**

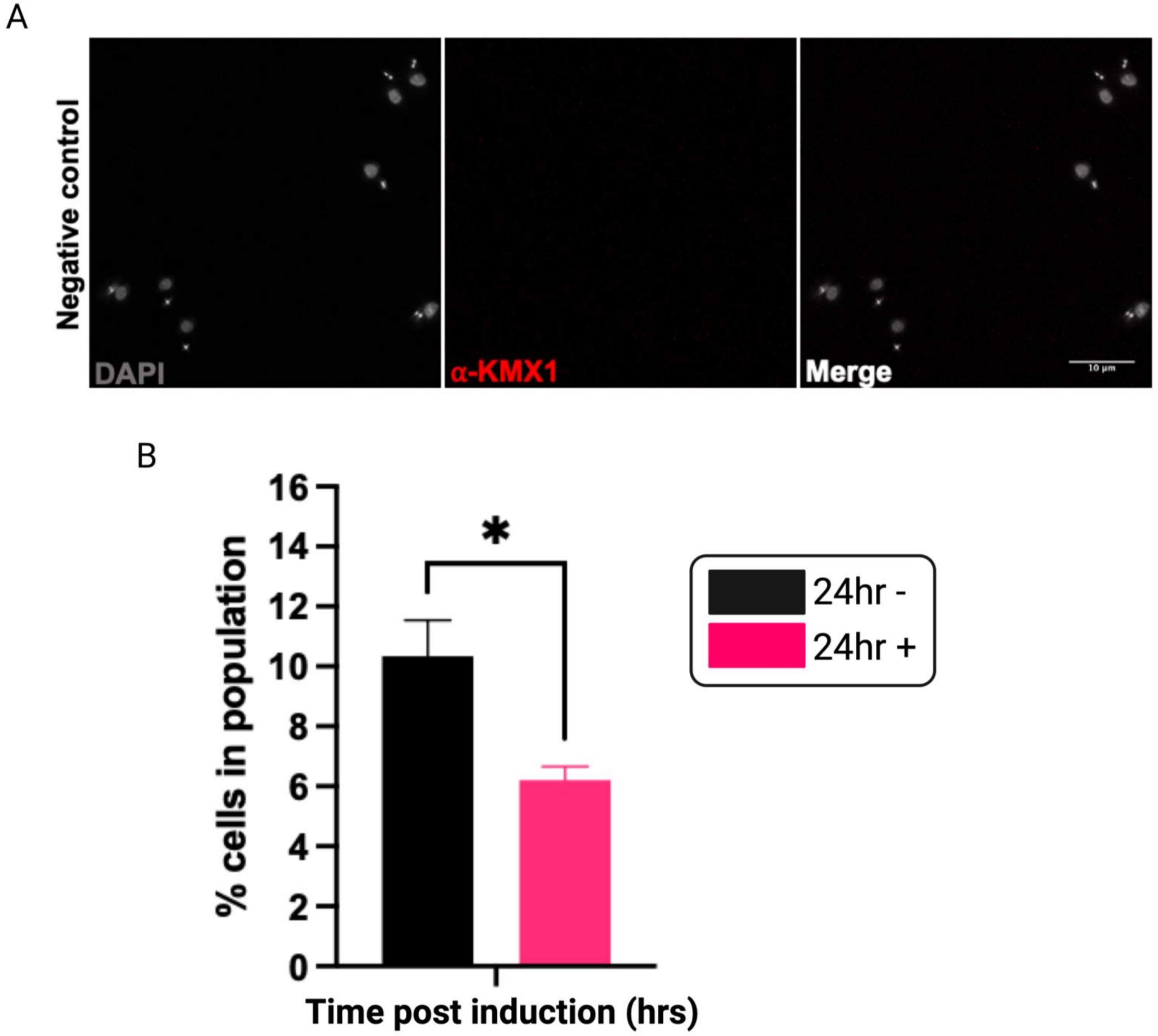
A
Negative control
DAPI
α-KMX1
Merge
10 µm
B
% cells in population
16
14
12
10
8
6
4
2
0
*
24hr -
24hr +
Time post induction (hrs)

**Figure S8**

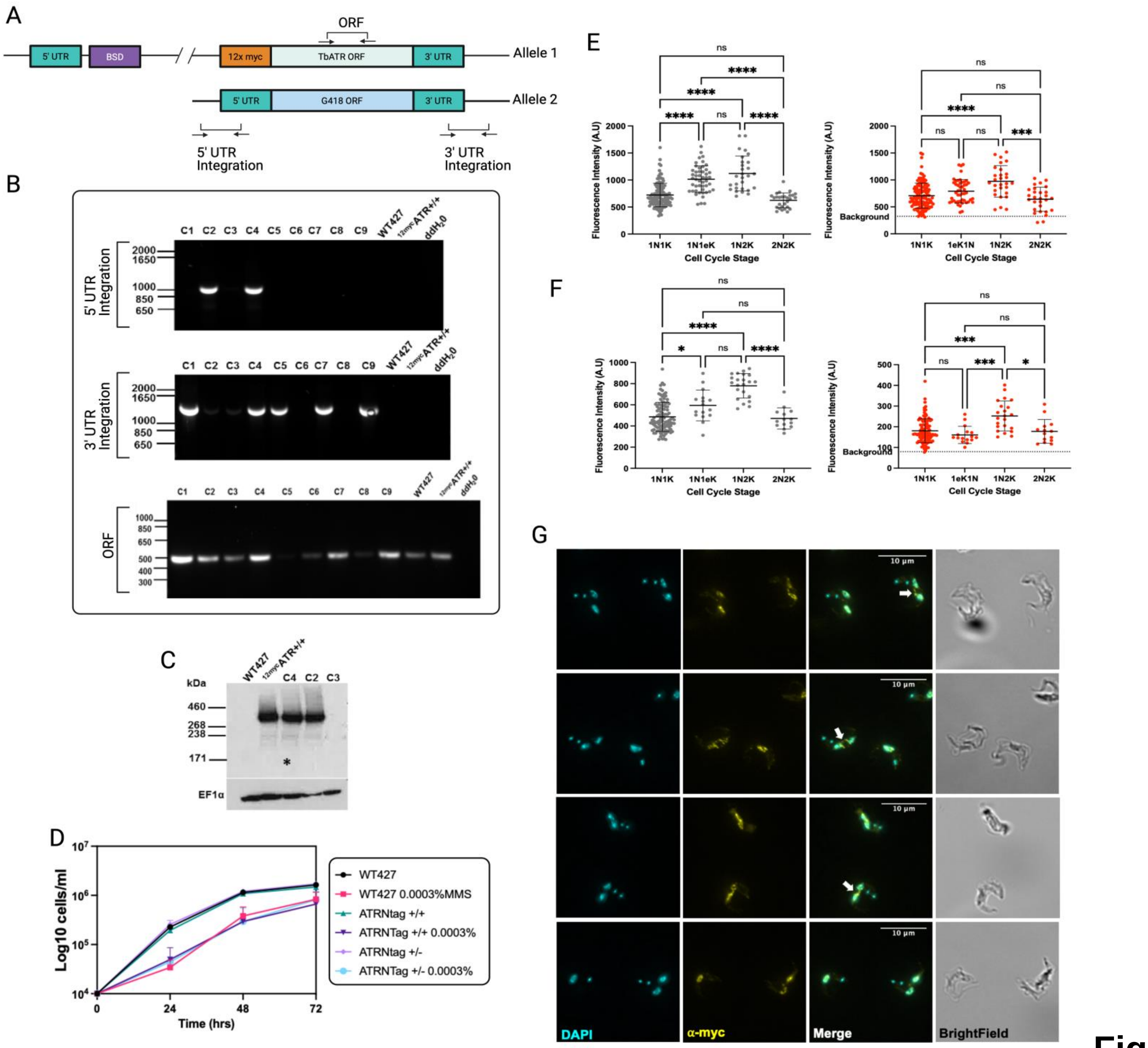
A
ORF
5' UTR
BSD
12x myc
TbATR ORF
3' UTR
Allele 1
5' UTR
G418 ORF
3' UTR
Allele 2
5' UTR Integration
3' UTR Integration
B
5' UTR Integration
3' UTR Integration
ORF
C
kDa
EF1α
D
Log10 cells/ml
Time (hrs)
WT427
WT427 0.0003%MMS
ATRNtag +/+
ATRNTag +/+ 0.0003%
ATRNtag +/-
ATRNTag +/- 0.0003%
E
F
Fluorescence Intensity (A.U)
Cell Cycle Stage
Background
G
DAPI
α-myc
Merge
BrightField

Figure S9
